# The Arabidopsis Hop1 homolog ASY1 mediates cross-over assurance and interference

**DOI:** 10.1101/2022.03.17.484635

**Authors:** Gaetan Pochon, Isabelle M Henry, Chao Yang, Niels Lory, Nadia Fernández- Jiménez, Franziska Böwer, Bingyan Hu, Lena Carstens, Helen T. Tsai, Monica Pradillo, Luca Comai, Arp Schnittger

## Abstract

The chromosome axis plays a crucial role in meiotic recombination. Here, we study the function of ASY1, the Arabidopsis homolog of the yeast chromosome axis associated component Hop1. Specifically, we characterized cross-over (CO) distribution in female and male meiosis by deep sequencing of the progeny of an allelic series of *asy1* mutants. Combining data from nearly 1000 individual plants, we find that reduced ASY1 activity leads to genomic instability and sometimes drastic genomic rearrangements. We further observed that COs are less frequent and appear in more distal chromosomal regions in plants with no or reduced ASY1 activity, consistent with previous analyses. However, our sequencing approach revealed that the reduction in CO number is not as dramatic as suggested by cytological analyses. Analysis of double mutants of *asy1* with mutants with three other CO factors, MUS81, MSH4 and MSH5 as well as the determination of foci number of the CO regulator MLH1 demonstrates that the majority of the COs in *asy1*, similar to the situation in the wildtype, largely belong to the class I, which are subject to interference. However, these COs are redistributed in *asy1* mutants and typically appear much closer than in the wildtype. Hence, ASY1 plays a key role in CO interference that spaces COs along a chromosome. Conversely, since a large proportion of chromosomes do not receive any CO, we conclude that CO assurance, the process that ensures the obligatory assignment of one CO per chromosome, is also affected in *asy1* mutants.

**Significant statement:** The regulation of the number and placement of cross-overs (COs) during meiosis is critical to ensure meiotic fidelity and promote new genetic combinations. Here, we investigated the function of one of the proteins of the chromosome axis, which plays a key role in CO formation: ASY1. Our results show that COs in *asy1* mutants are positioned closer to each other than in the wildtype and that, despite a roughly similar number of COs, not every chromosome receives a CO. With this, our results shed light on the mechanisms regulating two important but still poorly understood aspects of meiosis: CO assurance, which safeguards at least one CO per chromosome pair, and CO interference, which prevents two COs from occurring close to each other.

## Introduction

Meiotic cross-overs (CO) are critical to the generation of genetic diversity. Through COs, segments of homologous chromosomes (homologs) are exchanged during the first meiotic division, creating new and unique combinations of alleles. To this end, DNA double strand breaks (DSBs) are deliberately introduced into chromosomes in early prophase I. A fraction of these DSBs are then repaired by homologous recombination (HR) between the homologs, resulting in COs (1). HR relies on the resection of DSBs by the MRN complex generating 3’-overhanging single-stranded DNA, the search for sequence homology between the homologs, and subsequent strand invasion mediated by the RecA-related recombinases RADIATION SENSITIVE51 (RAD51) and DISRUPTED MEIOTIC cDNA (DMC1). Following DNA synthesis of the invading strand, and the capture of the second end of the initial DSB, a tetrahedral cross-strand intermediate is formed that is known as double Holliday junction (dHJ), which, when resolved, can result in a CO or a non-CO (NCO) (2, 3). NCOs can also be generated without the formation of dHJs (4).

COs are typically grouped into two classes, which can be distinguished by the molecular machinery that catalyzes the resolution of the dHJ. Type I COs rely on the ZMM group of proteins including the DNA mismatch repair proteins MUTL HOMOLOGUE1 (MLH1) (1). A hallmark of type I COs is that they experience positive interference, i.e., COs are more distantly spaced from each other than expected by chance (5). This results in a gamma-shaped distribution of COs across the genome. In contrast, type II COs are formed by a different pathway, which involves the endonuclease MMS4 AND UV SENSITIVE81 (MUS81) and are not subjected to interference (1).

Of key importance for meiotic recombination is the formation of the chromosome axis, by which a proteinaceous scaffold is formed along the DNA, and from which chromatin loops are thought to extrude (6). The chromosome axis plays a key role in DSB induction in many species and early steps of CO formation (7). Later in meiosis I, this axis becomes the lateral element in the synaptonemal complex (SC) that zips the two homologs together and promotes the maturation of COs in many species. An important, yet biochemically poorly understood protein, which is associated with the chromosome axis is Hop1 and its orthologs such as ASYNAPTIC1 (ASY1) in Arabidopsis. Earlier work has implicated ASY1/Hop1 in many central aspects of meiotic recombination, such as promoting interhomolog-biased DNA repair, chromosome pairing, SC formation, and CO production (8–14).

The number and pattern of CO formation varies between species, and can be modulated by environmental conditions (15). The number of COs is also different between male and female meiosis. In Arabidopsis, there are usually 10 to 12 COs formed during male meiosis, but typically only 6 to 7 during female meiosis (16). The majority of COs, i.e., approximately 85%, are type I COs, and the 2-3 COs per chromosome are typically evenly distributed along the chromosome in male meiocytes (17). In contrast, a single CO is typically observed per chromosome in female meiocytes, and it is more often located in the pericentromeric regions (16). Importantly, all chromosomes typically undergo at least one CO in both sexes. This is insured through a mechanism referred to as CO assurance, which has been detected in many species (18–20). How CO interference and assurance are brought about is currently not understood. In yeast, several mutants have been identified that affect CO interference, including topoisomerase II (TOPO II) (19). Whether these components function in a similar manner in other organisms remains unclear. One attractive model of how interference could be established is the “beam-film model” (21, 22). This model predicts that COs are formed at sites of high mechanical stress between the engaged homologs. Once a CO is formed, it locally releases this stress and this relaxation spreads along the chromosomes laterally, preventing the formation of a CO nearby (21, 22). However, how this tension stress and its relaxation are transmitted is not clear.

CO site selection appears to involve both epigenetic information, such as DNA methylation (23–26), and genetic factors such as the activity of the cell cycle kinase CYCLIN DEPENDENT KINASE A;1 (CDKA;1) (27, 28), as for instance shown in Arabidopsis. Interestingly, the number and position of COs has often been found to be modulated in polyploid plants, possibly to adapt to the challenge of faithfully assorting multiple chromosome sets during meiosis (29). Typically, polyploid species show a reduced CO number, and the remaining COs are often located at distal positions, a pattern which is also often found in diploid crop species such as maize (30). How this change in pattern is accomplished is not clear. It is also not clear whether allelic variants of the core meiotic machinery genes that appear to be associated with polyploidization, including alleles for *ASY1*, are functionally associated with this change in CO number and distribution (31).

Here, we have assessed the role of ASY1 in CO formation and distribution, using a genomics approach. To this end, we generated recombination maps of female and male meiosis for nearly 700 *asy1* mutant plants, and characterized their CO patterns in comparison to the WT. First, we observed many instances of full or partial chromosome aneuploidy in the progenies of the *asy1* mutants, confirming the importance of ASY1 for meiotic fidelity. Our work revealed that the overall number of COs is reduced in *asy1* mutants but this trend is less pronounced in the surviving progeny than previously estimated by cytological studies. We show that COs in *asy1* largely belong to the class I of COs and tend to closely localize at the telomeric and subtelomeric regions, highlighting a role for ASY1 in CO interference. Furthermore, the distribution of COs per chromosome is altered in plants with reduced ASY1 activities and a large percentage of chromosomes being devoid of any CO, suggesting that ASY1 is a crucial factor for CO assurance as well.

## Results

### Creation of mapping populations to study ASY1 function in meiotic recombination

To study the effect of ASY1 on meiotic recombination, we sought to dissect CO distribution through a backcross design of the two different *Arabidopsis thaliana* accessions Col-0 and L*er*-1. Therefore, we first generated an *asy1* null mutant using a CRISPR/Cas9 approach in the background of the accession Landsberg *erecta* (L*er*-1) (32). Cas9 was targeted to the *ASY1* coding region at the first exon-intron junction by a single guide RNA harboring 20 bases (Fig. S1A). A mutant allele (*asy1^Ler-1^*) harboring a guanine (G) insertion at the beginning of the second exon of *ASY1* was subsequently identified from the T2 generation that did not harbor the CRISPR/Cas9 T-DNA anymore. This G insertion results in a frameshift and premature stop codon after the translation of the first 13 amino acids. Homozygous *asy1^Ler-1^* mutants showed high levels of pollen mortality, and a high percentage of aborted seeds, matching the mutant phenotype of the previously identified *asy1* null mutant in the Columbia-0 accession (*asy1^Col-0^*) (Fig. S1B-E) (12, 33). That this allele represents a loss-of-function mutant was confirmed by the absence of ASY1 from male meiocytes by immuno-detection (Fig. S1F). A complementation test using the previously generated functional ASY1 reporter (*PRO_ASY1_:ASY1:GFP*) (34), showed that the fertility defects of *asy1^Ler-1^* mutants were fully rescued by this reporter, corroborating that the meiotic defects of this CRISPR/Cas9 induced allele are due to the loss of ASY1 function (Fig. S1B-E).

Next, the *asy1^Ler-1^* mutant was combined with either an *asy1^Col-0^* null mutant, or a previously generated hypomorphic *asy1* mutant, called *asy1^T142V^* (both in the Col-0 genetic background), in which the threonine residue of the presumptive CDKA;1 phosphorylation site at position 142 is replaced by the non-phosphorylatable amino acid valine (34). *In vitro*, the non-phosphorylatable mutant *asy1^T142V^* has reduced binding affinity to ASY3, a key component of the chromosome axis, and shows decreased chromosome association *in vivo* (*34, 35*). These combinations then resulted in the following Col-L*er* F1 hybrids: the first carries two null alleles of *ASY1* (referred to as **aa** from now on), and the second carries a null allele in L*er*-1 and the hypomorphic allele in Col-0 (*asy1^T142V-Col-0^ asy1^Ler-1^* referred to as **ta** from now on).

Finally, we produced 6 mapping populations by backcrossing these hybrids to WT Col-0 plants (Table 1): Specifically, the hybrids with reduced ASY1 activity were crossed as female parents to the Col-0 WT (referred to as **CC** from now on) to assess the effect of reduced ASY1 activity in female meiosis. These crosses are designated in the following as **aa x CC** (n=207) for crosses with the *asy1* null mutant and **ta x CC** (n=213) for crosses with the hypomorphic mutant *asy1^T142V^*. Reciprocal crosses were performed to monitor male meiosis, designated as **CC x aa** (n=216) for crosses with the *asy1* null mutant and **CC x ta** (n=214) for crosses with the hypomorphic mutant *asy1^T142V^*. As a control, we also generated F1 hybrids between the WT parental lines Col-0 and L*er*-1, and crossed them reciprocally with a Col-0 tester line giving rise to two additional populations, named **LC x CC** (n=117) and **CC x LC** (n=119). To characterize recombination in *asy1* mutants, the progeny of these six populations were subjected to whole-genome sequencing.

**Table 1.** Description of the crosses and number of diploid and non-diploid individuals from each cross type.

| Cross | Female parent | Male parent | N (euploid) | N and (%) non-diploids | Total N |
| --- | --- | --- | --- | --- | --- |
| LC x CC | Col-0 / Ler-1 | Col-0 | 117 | 0 (0.0%) | 117 |
| ta x CC | *asy1*<sup>T142V-Col-0</sup> / *asy1*<sup>Ler-1</sup> | Col-0 | 146 | 67 (31.5%) | 213 |
| aa x CC | *asy1*<sup>Col-0</sup> / *asy1*<sup>Ler-1</sup> | Col-0 | 176 | 31 (15.0%) | 207 |
| CC x LC | Col-0 | Col-0 / Ler-1 | 119 | 0 (0.0%) | 119 |
| CC x ta | Col-0 | *asy1*<sup>T142V-Col-0</sup> / *asy1*<sup>Ler-1</sup> | 155 | 59 (27.6%) | 214 |
| CC x aa | Col-0 | *asy1*<sup>Col-0</sup> / *asy1*<sup>Ler-1</sup> | 200 | 16 (7.4%) | 216 |

C: Col-0, t: *asy1*<sup>T142V-Col-0</sup>, a: *asy1*<sup>Ler-1</sup> or *asy1*<sup>Col-0</sup> null alleles.

### Reduction of ASY1 activity results in aneuploid offspring and genome instability

Low recombination frequencies often cause the formation of univalents and hence unequal chromosome distributions and aneuploidy in the resulting spores (11). Previous cytological data (12, 33, 35, 36) suggested that CO numbers might be reduced in *asy1* mutants since not all homologs are able to form bivalents. Consistent with these earlier studies, we also found univalents in meiotic chromosome spread analyses of male meiocytes produced by ta plants (Fig. S2). To investigate the effect of these univalents on the karyotype of the resulting progenies, we screened each of these populations for the presence of aneuploidy. Looking for variation in sequencing read coverage across the whole genome, we found aneuploid plants in the progenies of all four crosses involving *asy1* mutants, while none were found in the control WT crosses (Table 2 and Fig. 1).

**Fig. 1.**
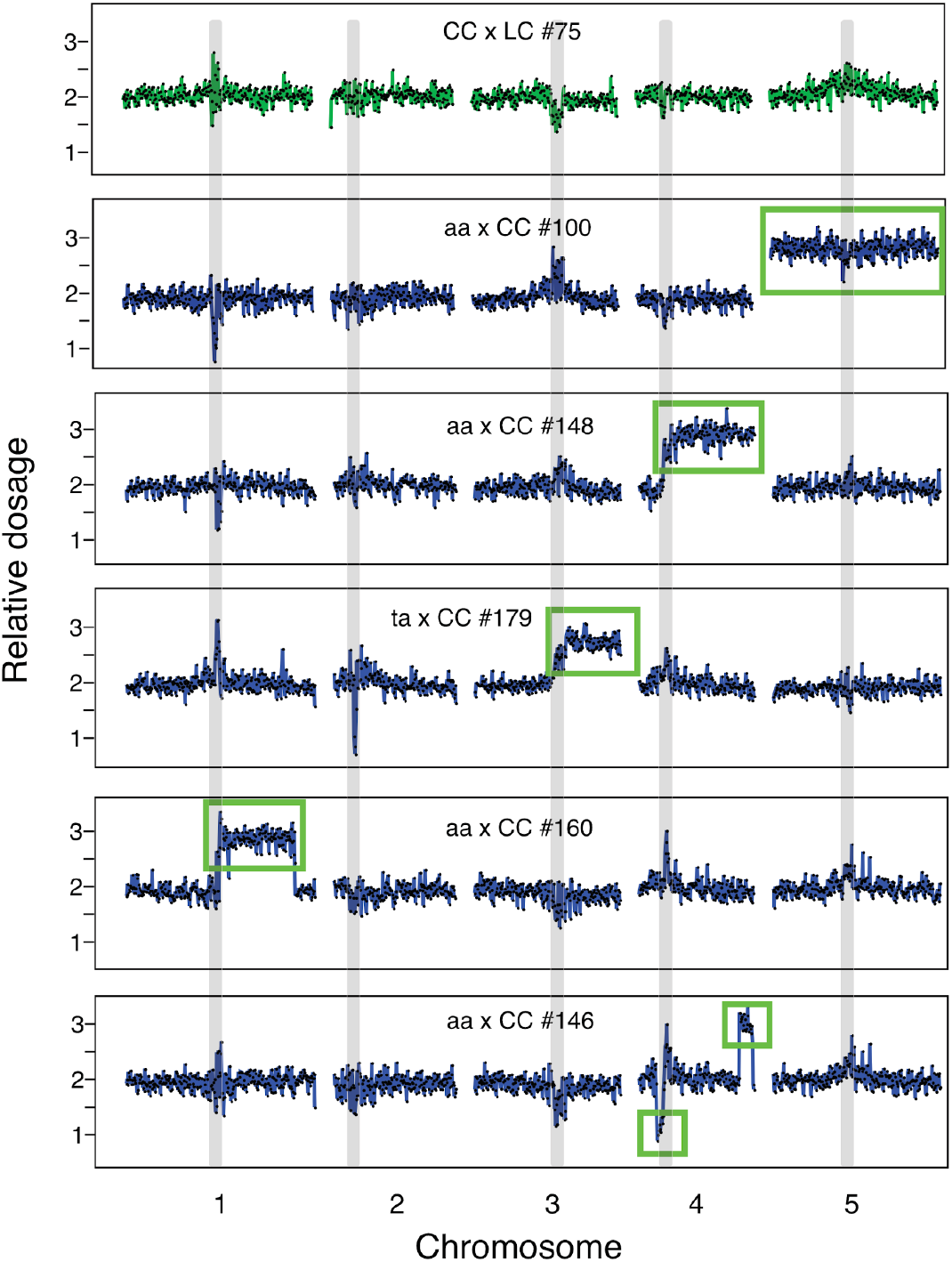
Aneuploidy in the progeny of the *asy1* mutants. Dosage variation in six representative individuals from different crosses (CC x LC #75, aa x LC #100, aa x CC #148, ta x CC #179, aa x CC #160, and aa x CC #146). Each panel represents one individual and, for each individual, the five chromosomes are shown. For each individual, relative dosage, e.g., ∼ 2 for normal diploids, is depicted based on short-read sequence coverage variation after being normalized to the whole population. Dots represent consecutive non-overlapping 100kb bins. None of progeny of wild-type control crosses (CC x LC and LC x CC) exhibited aneuploidy (top panel, shown in green). Crosses involving *asy1* or *asy1^T142V^* mutants produces many aneuploid progeny (panel 2-6, colored in blue), including full trisomics (panel 2, aa x LC #100), partial trisomics (3rd panel, aa x CC #148), or more complex aneuploids (panels 5 and 6, aa x CC #160 and aa x CC #146). Many of the progeny of the crosses involving the hypomorphic *asy1^T142V^* mutant carried an extra copy of the bottom arm of chromosome 3 (Chr. 3B) (Table 2), suggesting that it originated from the parental line (panel 4, ta x CC #179). The regions that are present in additional or missing copies are indicated by the green rectangles. Centromeric and pericentromeric regions (grey vertical bands) consistently exhibited variation that mimicked dosage variation, but more likely originated from allelic variation between Col-0 and L*er*-1 in these repeat-rich regions. These variations were consistently observed in all populations and were not recorded as aneuploidy.

**Table 2.**
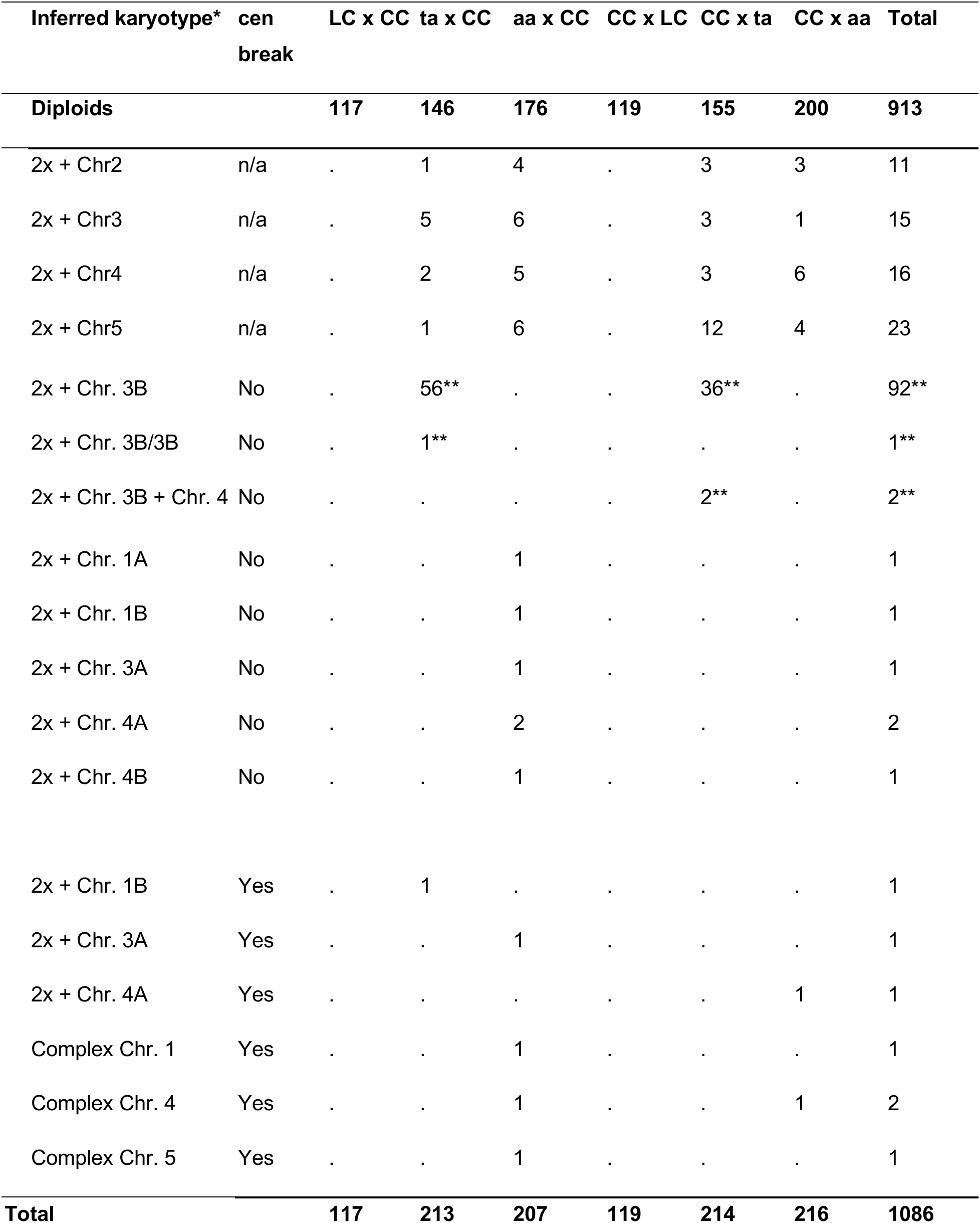
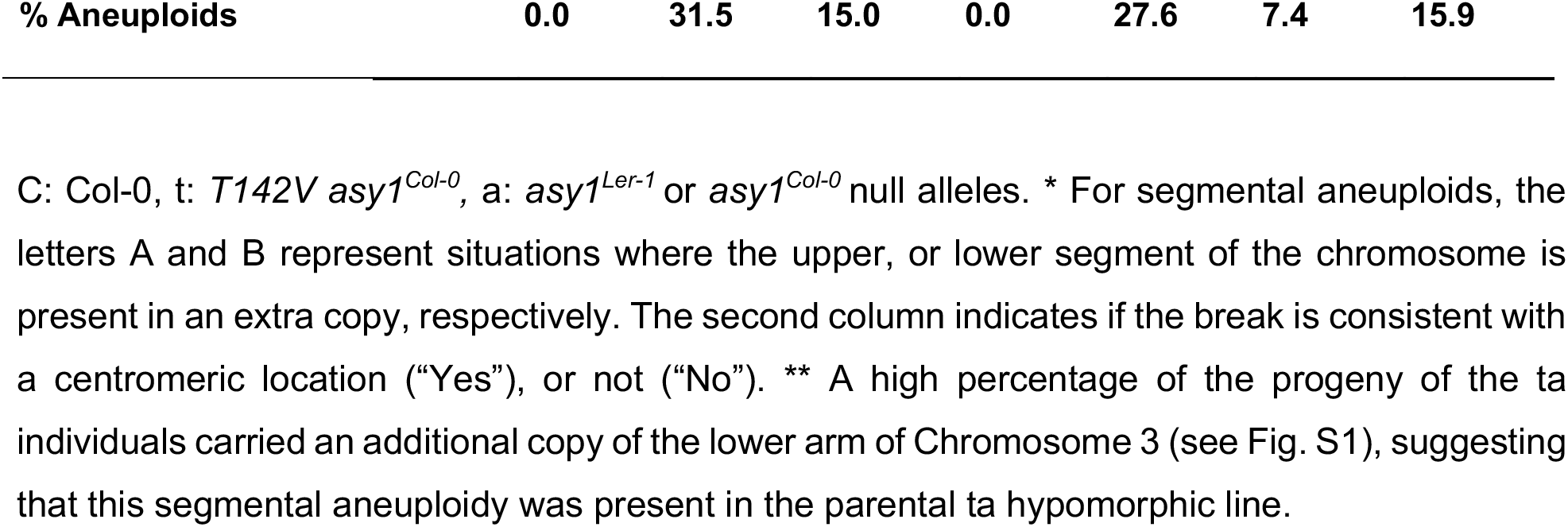
Number of diploid and aneuploid individuals obtained from each cross type.

Most of the aneuploids recovered were trisomics, carrying one additional copy of a single chromosome (Fig. 1, second panel). Trisomies of all chromosome types were recovered, except for chromosome 1 (Table 2), perhaps because of more severe selection against gametes carrying additional copies of this particular chromosome, as observed previously (37, 38), or because of the more severe phenotypic consequences associated with additional copies of chromosome 1 in *A. thaliana* (39). Conversely, we did not detect any instance of a missing chromosome, probably because it is lethal or severely detrimental at the gametic stage, consistent with previous work (40–42).

Interestingly, we also found examples of segmental aneuploids, carrying only a part of an additional chromosome (Fig. 1, panels 3-6 and Table 2). The karyotypes of some of segmental aneuploids could have originated from a single miss-repaired DNA DSB (Fig. 1, panels 3, 4), in which case only a terminal segment remains. In addition, we find individuals carrying signs of two or more breaks (Fig. 1, panels 5, 6), resulting in more drastically rearranged genomes. Comparing the approximate location of these breaks with the expected position of the centromeres (The Arabidopsis Genome Initiative), suggested that more than half (9/17) of these miss-repaired breaks occurred at or around the centromeres (Table 2, second column). While this pattern possibly suggests a direct or indirect role of ASY1 in the formation and/or repair of DSB near centromeres, breaks at centromeres are also often enriched after gamma radiation (41), in the progeny of aneuploids (38), and after genome elimination (40), suggesting that centromeric regions are in particular sensitive to DNA stresses.

Most of the individuals exhibiting segmental aneuploidy originated from crosses involving the *asy1* mutants as a female (Table 2). This overrepresentation in females is consistent with the idea that segmental aneuploids are less fit than euploids, and hence are counter selected during pollen tube growth. Moreover, segmental aneuploids appeared much more frequently in crosses involving *asy1* null mutants, as we only found one case originating from a cross with the hypomorphic mutant (Table 2), suggesting that genomic instability scales with the level of ASY1 reduction.

Strikingly, one individual (#102) from the aa x CC cross exhibited signs of high levels of genomic instability (Fig. S3). Plant #102 carried an extra copy of the top arm of chromosome 1 but coverage information for that region of the genome was highly variable compared to the diploid individual from the same cross (Fig. S3, panel 1, 3). Comparison with other segmental aneuploids of chromosome 1 confirmed that this level of variability did not originate from methodological artifacts associated with the presence of a third copy of the corresponding sequences (Fig. S3, panel 2). Such a catastrophic genome restructuring event is reminiscent of chromothripsis, when laggard chromosomes are incorporated into micronuclei, and undergo massive DSBs, resulting in restructuring and copy number variation clustered on a single chromosome (43). Chromothripsis is often associated with cancer in humans (43), but has also been observed in *A. thaliana* following CENH3-mediated genome elimination (40). The extent of dosage variation observed here is very high, with no clear and consistent copy number shift, suggesting that copy number varies so often that it cannot be captured within our 100 kb bins, or that the tissue sampled was chimeric, and that we are witnessing a mixture of copy number states, or both. It is likely that other plants carrying severely rearranged chromosomes were strongly counter selected and are not found in our experiments. Thus, this sample provides a unique opportunity to witness a snapshot of the extent of meiotic instability of *asy1* mutants.

Unexpectedly, both of the crosses involving the hypomorphic mutant produced a population of plants enriched in segmental trisomy for chromosome 3B (Fig. 1, individual ta x CC #179). The fact that this particular karyotype was clearly overrepresented in those two crosses (56/213 for ta x CC and 36/214 for CC x ta), but never found in the other four crosses (Table 2), suggests that the hybrid plant carrying the hypomorphic *asy1* mutation itself carried this segmental aneuploidy, and transmitted it to a subset of its progeny. Possibly, this additional copy of chromosome 3B originated from defective meiosis in the parental *asy1^T142V^* or *asy1^ler-1^* mutants that were used to produce the F1 hypomorphic ta mutant (see Methods). Since the additional copy of chromosomal segment would strongly interfere with the identification of COs on this particular chromatid using the method applied in this study, we discarded all aneuploid progenies from further analysis. The final number of diploid individuals remaining for each population ranged between 117 and 200 (Table 2).

### Number and position of COs are affected by reduced activity of ASY1

To characterize the number and position of COs in the WT and *asy1* mutant lines, each remaining diploid individual was genotyped at each of 239 non-overlapping 500kb marker bins covering the entire *A. thaliana* genome. The data were input into r-qtl (see Methods) and separate genetic maps were created for the six populations (Fig. S4). In all cases, the marker order was consistent with the physical order of the bins. We next identified the position of all COs detectable in these samples (Supplemental Dataset 1).

The mean number of COs per chromosome varied depending on whether the *asy1* mutant was the male or the female, the chromosome type, and the level of ASY1 activity (Fig. 2). Specifically, the number of COs per chromosome was not significantly different for the control or mutant crosses for the subtelocentric chromosomes (chromosomes 2 and 4). On the other hand, there was a significant decrease in CO number for chromosomes 1, 3 and 5 on the male side, and chromosomes 1 and 3 on the female side (t-test p-values < 0.05 or less, Fig. 3). In most of these cases, the hypomorphic allele exhibited CO numbers that were intermediate between the control crosses and the crosses involving the *asy1* null allele.

**Fig. 2.**
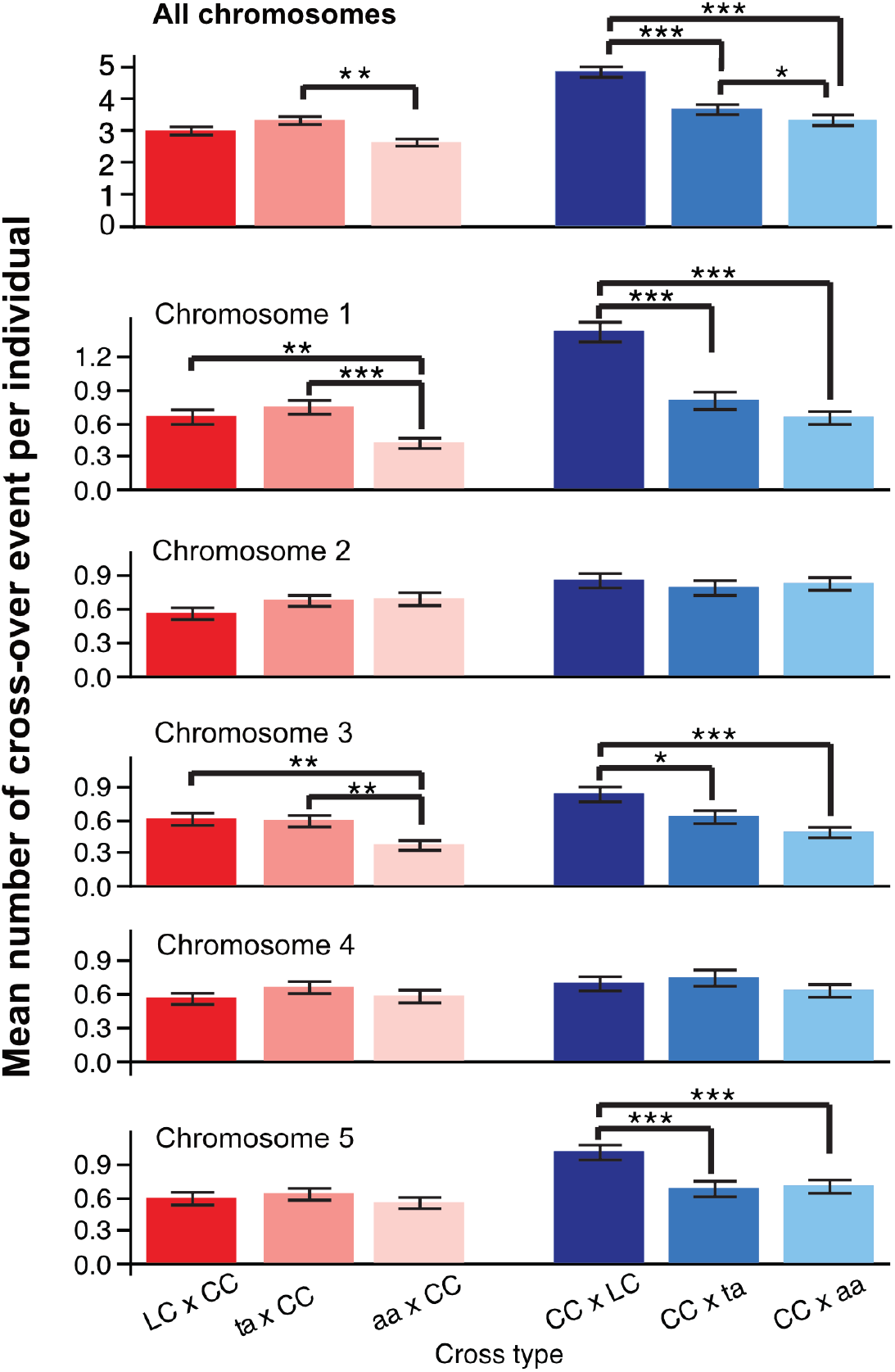
Mean number of CO per individual. For each cross-type, the mean number of CO events was calculated for each individual (top panel) and for each chromosome and each individual (all other panels). CO events were only detectable for the hybrid (L*er*-Col) parent, not the Col-0 parent, and therefore represent either events that occurred on the paternal side (blue bars), or the maternal side (red/pink bars) only. Since each meiosis produces four gametic cells and each CO event can only be detected from two of four gametes, the average number of CO per chromosome per individual indicates only half of the CO events formed in each meiosis. For each cross type and chromosome, the mean and standard errors are represented. Within a cross type and a cross direction, significant differences between the means were tested on a pair-wise basis using Student’s t-test. Significant comparisons are indicated as follows: ***: p-value < 0.0001, **: p-value < 0.01, *: p-value < 0.05.

**Fig. 3.**
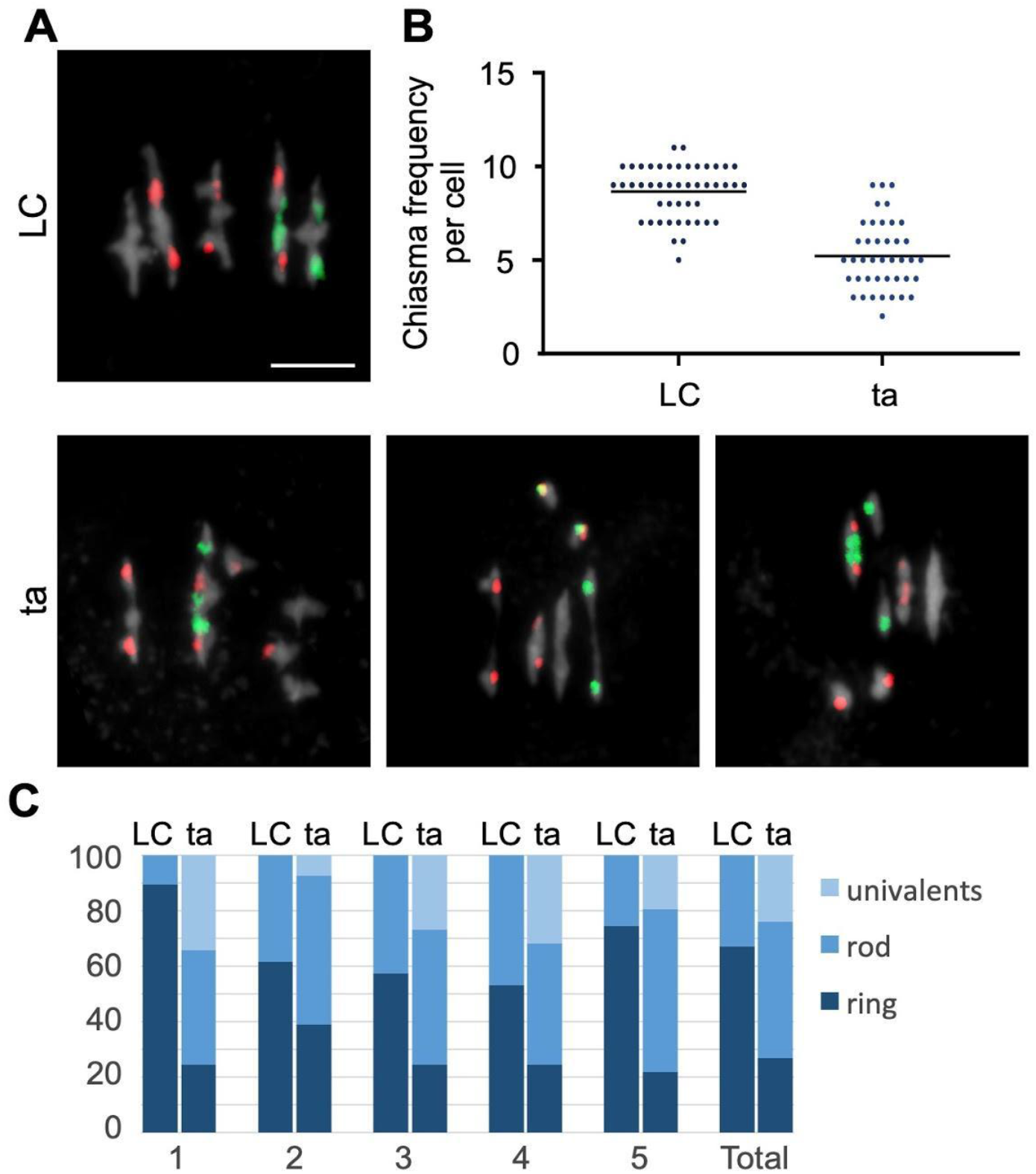
Determination of chiasmata number in *asy1* mutants using FISH. (A) Representative metaphase I nuclei of LC (up) and ta (down) meiocytes using FISH to identify the different chromosome types. Red signals (5S rDNA loci) are present in chromosomes 3, 4 and 5, whereas green signals (45S rDNA loci, NORs) appear in the acrocentric chromosomes 2 and 4. DAPI staining appears in grey. Examples of ta cells include univalents in different chromosomes: 1 and 3 (left); 4 (middle); 5 (right). Bars represent 5 mm. (B) Quantification of chiasmata in metaphase I cells. Each dot represents an individual cell and bars indicate the mean. (C) Percentage of pairs of univalents, rod bivalents (with chiasmata only in one chromosome arm), and ring bivalents (with at least one chiasma on both chromosome arms) in the wildtype LC hybrids (left) and the ta hybrids (right).

Next, we analyzed the chiasma formation in *asy1* mutants by chromosome spreads of male meiocytes at metaphase I. This analysis revealed a significant decrease in the chiasma frequency of *asy1* mutants (*asy1^Ler-1^*: 2.62 ± 1.39, n=47; *asy1^Col-0^*: 2.78 ± 1.17, n=65; *asy1^T142V^*: 5.2 ± 0.3, n=41 vs. 8.7 ± 0.2 in the WT, n=47). To have a resolution at chromosomal level, FISH experiments were performed to distinguish different chromosomes using the 5S and 45S rDNA probes in ta mutants in which the occurrence of aneuploids is not so severe as that in *asy1* null mutants. We found that all chromosomes, except for chromosome 2, showed a reduction in CO formation (Fig. 3 and table 3), and this reduction was higher for the large chromosomes 1 and 5, consistent with earlier work analyzing *asy1* null mutants (12, 33) (Fig. 3C).

**Table 3.** Mean chiasma frequencies per chromosome.

| Genotype |  | Chiasma frequency per chromosome |  |  |  |  | Total |
| --- | --- | --- | --- | --- | --- | --- | --- |
|  |  | 1 | 2 | 3 | 4 | 5 |  |
| LC |  | 2.1 | 1.6 | 1.6 | 1.5 | 1.8 | 8.7 |
|  |  | 24.7% | 18.6% | 18.3% | 19.5% | 20.7% |  |
| ta |  | 0.9 | 1.4 | 1.0. | 1.0 | 1.0 | 5.2 |
|  |  | 17.2% | 27.4% | 18.1% | 17.4% | 20.0% |  |
| Student's t-test | *t* | 8.12 | 1.92 | 4.56 | 4.11 | 6.19 | 9.94 |
|  | *p* | 0.000 | 0.058 | 0.000 | 0.000 | 0.000 | 0.000 |

The percentages represent the chromosome contribution to the total chiasma frequency. We found differences for all five chromosomes, except for chromosome 2. The reduction was higher for the large chromosomes 1 and 5.

To examine whether the reduction in bivalent formation in *asy1* mutants is due to a reduction in the formation of DSBs, we analyzed RAD51 foci in leptotene/zygotene. We found that RAD51 foci numbers in male meiosis of aa (136.8 ± 10.9; n = 14) and ta (134.1 ± 17.6, n=15) were not significantly different from those of the WT LC hybrids (134.6 ± 14.2, n=12; aa vs LC p=0.89; ta vs LC p=0.99) (Fig. S5). Therefore, the observed reduction in CO number in the *asy1* hybrids does not originate from reduced DSB formation, consistent with previous analysis of *asy1* null mutants (9).

The discrepancy between the observed chiasmata and the COs identified by sequencing, especially for chromosome 4, for which we did not find any significant reduction in CO number in our genetic maps, prompted us to analyze CO distribution within each chromosome (Fig. 4 and Fig. S6). For ease of visualization and comparison, we expressed CO position as the distance to the telomeres, expressed in 0.5 Mb units, and are presenting data from the left and right arms separately, to distinguish the effect of the mutations from those of the short arm of the subtelocentric chromosomes. The distribution of COs was markedly different for the ta and aa crosses, compared to the control crosses, in both cross directions (Fig. 4A). Specifically, COs were inclined to localize near the telomeres in the *asy1* mutant crosses, while they were more evenly distributed along the length of the chromosomes in the WT control crosses. The hypomorphic mutant exhibited an intermediate behavior. This trend was not only visible when data for all chromosomes were pooled (Fig. 4A), but also for each individual chromosome type (Fig. 4B), with the exception of the left arms of chromosomes 2 and 4, which are too short to detect changes in distribution. This telomere-proximal localization of COs is consistent with previous and our cytological analyses of spread chromosomes, which revealed long-stretched bivalents due to the chiasmata in chromosomal arm ends (Fig. 3) (12, 33). These findings also resolved the apparent difference to the cytological work that does not have the resolution to reveal closely spaced COs.

**Fig. 4.**
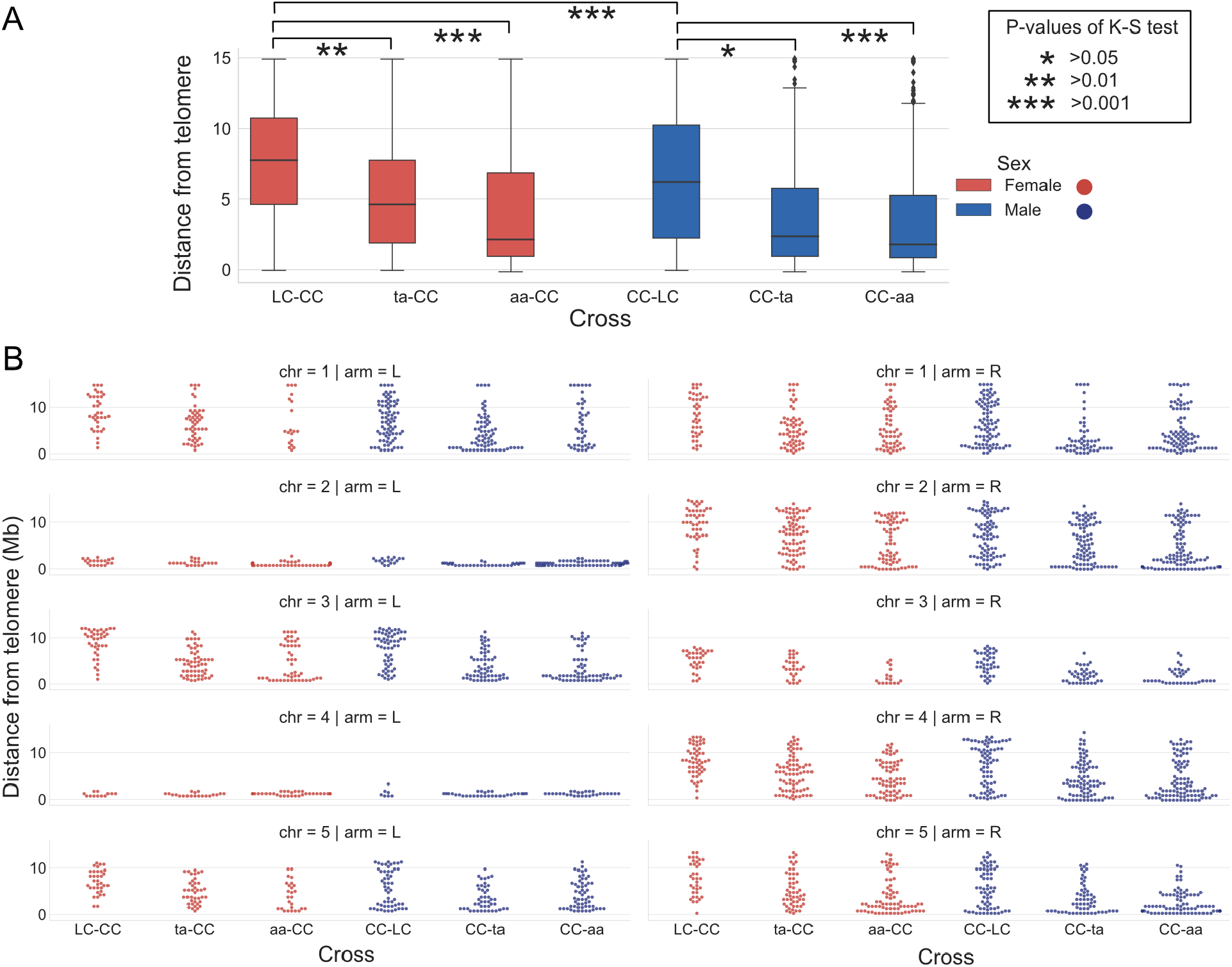
Distribution of CO position in the different crosses. **A.** Box plot of the cumulative distributions of CO-telomere distance. Significance of comparisons was tested by the Kolmogorov-Smirnov test (K-S) **B.** Swarm plots of CO-telomere distance per chromosome. Distance from CO to telomeres is shown and the maximum distance for each arm is at the centromere.

### CO interference is reduced in *asy1* mutants

The observed overaccumulation of COs in distal chromosome positions could be due to two different mechanisms. On the one hand, the number of interference-sensitive type I COs could be reduced at the benefit of additional type II COs, which are not subject to interference. On the other hand, type I COs, which are sensitive to interference, might lose their sensitivity in meiocytes with reduced or no ASY1 activity.

To discriminate between these two possibilities, we generated double mutants of *asy1^Col-0^* with mutants in *MSH4* (also in Col-0 background), one key component of type I CO formation, and double mutants of *asy1* with *mus81* (in Col-0 as well) in which type II COs are reduced. Depleting *MSH4* in *asy1* caused a drastic reduction of chiasma numbers, i.e., in 25 meiocytes of the *asy1 msh4* double mutants a chiasma frequency of only 0.07 CO per meiocyte was observed while this number, as shown above, is 2.78 ± 1.17 (n=65) in *asy1* single mutants.

Thus, the majority of the remaining COs in *asy1* belong to the type I class of COs. Consistently, the *asy1 mus81* double mutants did not show a strong reduction of chiasmata i.e., from 57 meiocytes, a chiasmata frequency of 1.9 ± 0.75 per meiocyte was obtained, suggesting that type II COs are present only at a small frequency in *asy1* mutants. Thus, the loss of ASY1 does apparently not grossly alter the relative proportion of type I to type II COs found in the wildtype.

To complement the genetic analysis, we performed immunolocalization analysis of MLH1, which marks type I COs, in male meiocytes (Fig. 6A,B). In WT LC hybrids, we observed an average of 7.2 ± 0.3 MLH1 foci per meiocyte (n=48), matching previous analyses. Strikingly, we did not find significant changes in the frequency of MLH1 foci in the aa plants (7.8 ± 0.3, n=35; p=0.406). A similar number was also found in the ta plants (7.8 ± 0.3, n=52; p=0.405). The slight discrepancy between the determination of COs in *asy1* mutants by sequencing with the number of MLH1 foci possibly indicates that not all MLH1 foci maturate into COs. However, although the MLH1 antibody and scoring of MLH1 foci has been often used in the past, we could fully exclude technical difficulties and for instance sometimes chromosomes are not clearly stained making it difficult to evaluate the results of the immuno-detection.

Therefore, we sought to complement the MLH1 localization by a previously established live cell imaging setup of meiocytes. To achieve this, we first generated a genomic reporter of MLH1 in which the GFP fluorescent tag was inserted immediately before the stop codon (*PRO_MLH1_:MLH1:GFP*, called *MLH1:GFP*), and then transformed this construct into *mlh1* mutants. In contrast to the severe fertility reduction of *mlh1* mutant plants, mutants harboring the *MLH1:GFP* construct were almost fully rescued as shown by the fully developed silique and good pollen viability, suggesting that MLH1:GFP is functional (Fig. S7A-C).

Next, the expression and localization of MLH:GFP at different stages of male meiosis was imaged by laser scanning confocal microscopy. We found that at pre-meiosis, i.e., S-phase, MLH1-GFP was expressed in entire meiocytes with many bright foci in the nuclei presumably highlighting DNA mismatch repair sites, which arise during replication (Fig. S4D). Subsequently, from early to mid prophase I, MLH1:GFP was only diffusely present in both the cytosol and nuclei of male meiocytes, and no particular pattern was observed (Fig. S4D). However, when meiocytes reached late prophase I, e.g., late pachytene or diplotene stages, clear foci appeared in the nuclei, presumably marking type I CO intermediates (Fig. S4D).

Next, to check CO distribution, *MLH1:GFP* reporter was introgressed into *asy1* mutant plants and compared to wild-type plants bearing this construct. To cover all MLH1 foci in a meicocyte, z-stacks with 0.8 ìm spacing intervals were acquired (Movie S1-4), and the foci were counted using the image analysis software Fiji. Consistent with our immunodetection (Fig. 6B), we found that there was a similar number of MLH1 foci in *asy1* mutant (9.13 ± 1.52, n=37) when compared to the wildtype (9.42 ± 1.36, n=34), corroborating that type I COs are formed in the mutant, at least when recombination progressed to this point, at a normal level. Notably, compared to the wildtype, very closely located MLH1 foci were frequently observed in *asy1* mutants (Fig. 6C, Movie S1-4).

The preference of telomere-proximal formation of CO in *asy1* mutant, together with the observation that these COs are of type I with largely the same number of MLH1-positive foci suggest that ASY1 might play a role in CO interference. To further investigate this possibility, we compared the distribution of distances between adjacent COs in the 6 different crosses (Fig. 5A-C). Inter-CO distance followed a broad distribution that appeared skewed toward short values for the ta and aa plants. Both female and male distributions in the aa mutant were significantly different from that of the corresponding WT crosses (Fig. 5A and B, Kolmogorov-Smirnov adjusted for multiple testing p=0.03, 0.02, respectively). To further probe this skewness, we divided inter-COs segments into two classes, close and far, arbitrarily changing the dividing line between these two classes and observing the effect on the results (Fig. 5B and C). The female control only yielded few instances of double COs (Fig. 5B, LC-CC cross), restricting our ability to observe significant differences between cross types. The ta-CC cross was significantly enriched in distant double COs (Fig. 5A and B). Nevertheless, significant enrichment for close events in female meiosis was demonstrated in the aa-CC cross when close and far bins were divided at 10 Mb (Fig. 5C, 10 Mb units). The higher recombination frequency in male meiosis provided more cases of syntenic COs than in female meiosis. Thus, the male CC-aa cross proved significant for any size bin and the male ta mutant displayed strong and significant differences above 1.255 Mb (Fig. 5C).

**Fig. 5.**
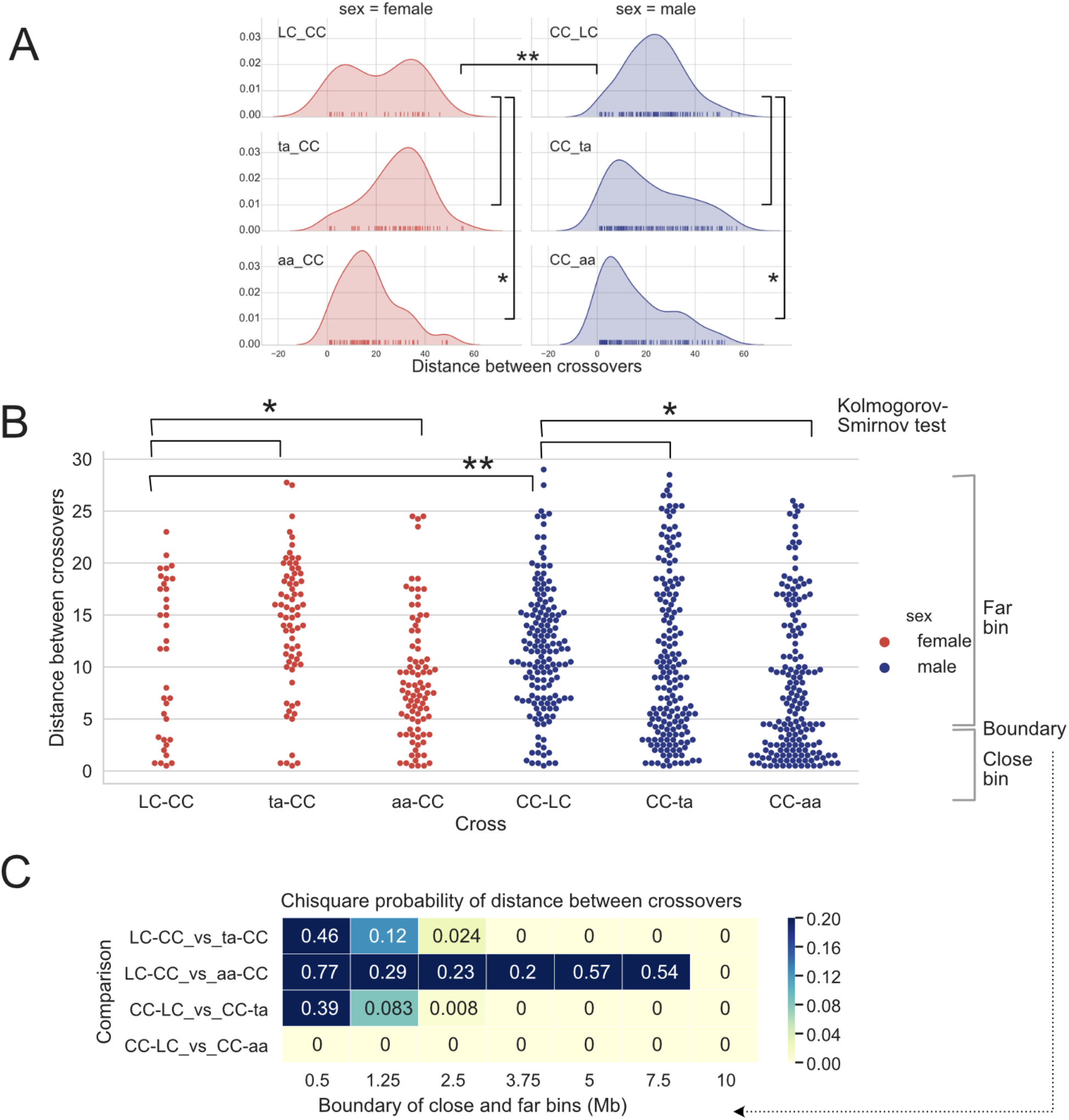
Frequency of COs and distance between Cos. CO to CO distance distribution displayed as (**A**) Kernel Density Estimation and (**B**) swarm plots, using in 1 Mb units. Knock out of *ASY1* is associated with reduced distance between COs. **C.** Chi-square probability for comparison of close vs. far binned CO-CO distances. The respective upper limit of the close bin varies and is shown below each column of p values. Yellow denotes significant difference by the Chi square test. Note that significance for the ta-CC cross was caused by an increase of the distant CO class, as opposed to an increase in the number of close COs, as observed in the aa crosses. Significance thresholds: * <0.05, ** <0.01.

A direct measurement of interference such as those obtained by the classical rate of observed vs. expected double COs, was not applicable because the number of individuals used for each cross limited the number of COs and statistical power. Nonetheless, measuring interference on a sliding, non-overlapping window of 4.5 Mb, suggested reduction of interference at multiple genomic intervals for both mutants, the aa null mutation displaying stronger reduction than the ta mutation (Fig. S8). The overall distribution of interference values for WT control crosses and *asy1* mutant crosses was highly significantly different (Fig. S8, Kolmogorov-Smirnov test adjusted for multiple testing p< 0.001 for all comparisons).

### CO assurance is affected in the ASY1 mutants

The above data indicated that ASY1 is required for proper number and distribution of COs along chromosomes. However, a telomeric or subtelomeric CO is sufficient to keep bivalents together, preventing a random distribution of homologs in meiosis I, and subsequent aneuploidy, as seen for instance in many crop species in which COs are usually distally located (30, 44, 45). Likewise, a low number of COs, does not necessarily lead to the formation of univalents as exemplified by female meiosis in Arabidopsis during which only about 6 to 7 COs are formed but every chromosome receives at least one CO (16). Thus, according to our sequencing data and chiasma counting (Fig. 2 and Table S2), the 5 to 6 COs formed in female meiosis and especially the 6 to 7 COs (reflecting the situation in female WT meiosis) formed in the male meiosis of *asy1* null and hypomorphic mutants could be roughly sufficient to equip every chromosome with a CO obeying CO assurance. However, *asy1* mutants show a high degree of univalent formation (Fig. 3C, Table S1), as also reported previously (12, 33). Consistently, we also found significantly more non-recombined chromosomes in our *asy1* backcross populations compared to the WT control cross (Fig. 7 and Table S3). This suggested impaired CO assurance as well.

**Fig. 6.**
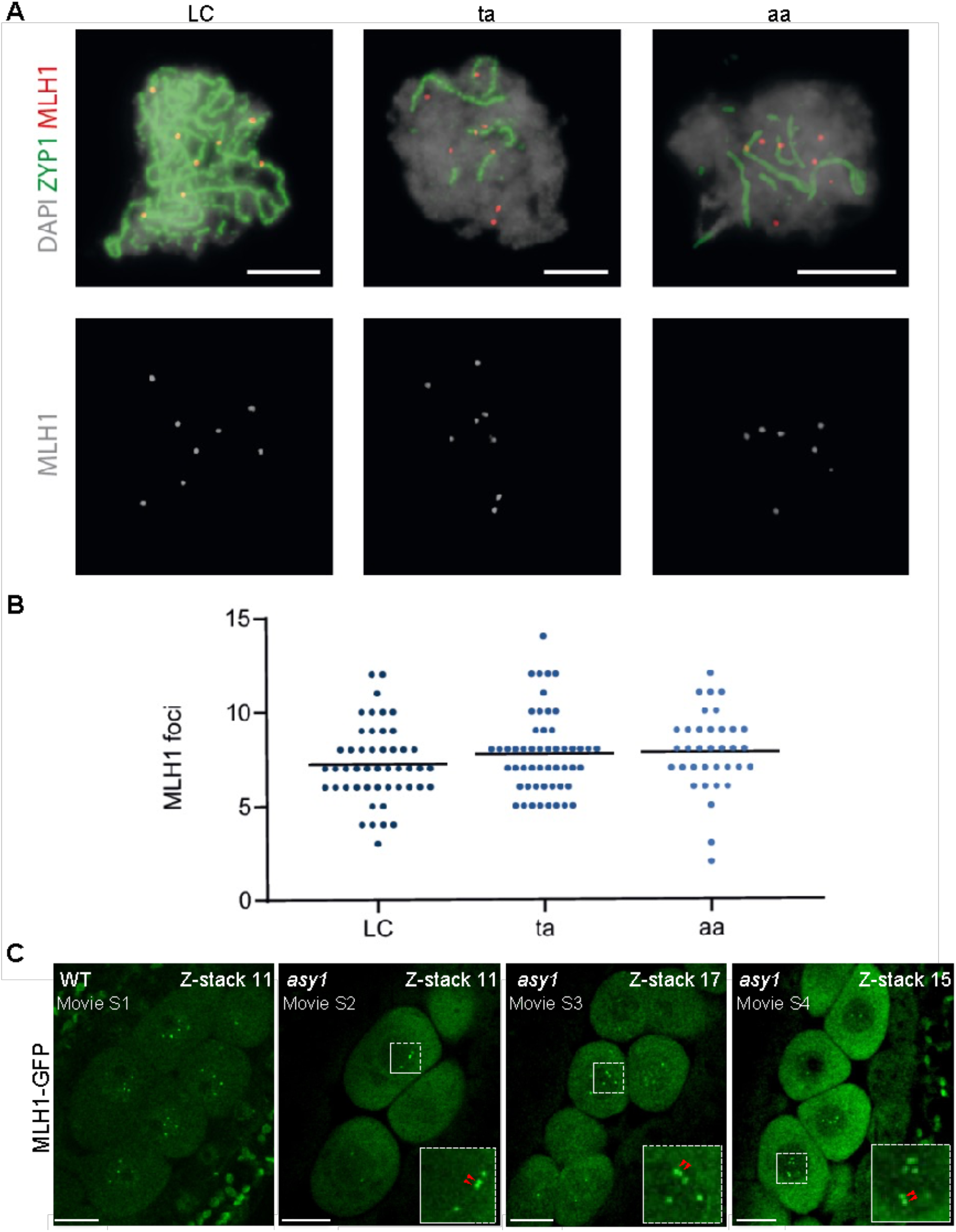
Analysis of MLH1 foci. (A) Representative images showing pachytene (LC) and pachytene-like (ta, aa) meiocytes after immunolabeling for detecting the class I CO marker MLH1 (red). The central element protein of the SC ZYP1 is detected in green and DAPI staining appears in grey. Bars represent 5µm. (B) Comparison of the number of MLH1 foci. Each dot represents an individual cell and bars indicate the mean. (C) Presence of MLH1 foci in wildtype (WT) and *asy1* mutant plants. One of the z-stacks were shown from Movie S1-4. Red arrows indicate the closely located MLH1 foci. Bars represent 10µm.

**Fig. 7.**
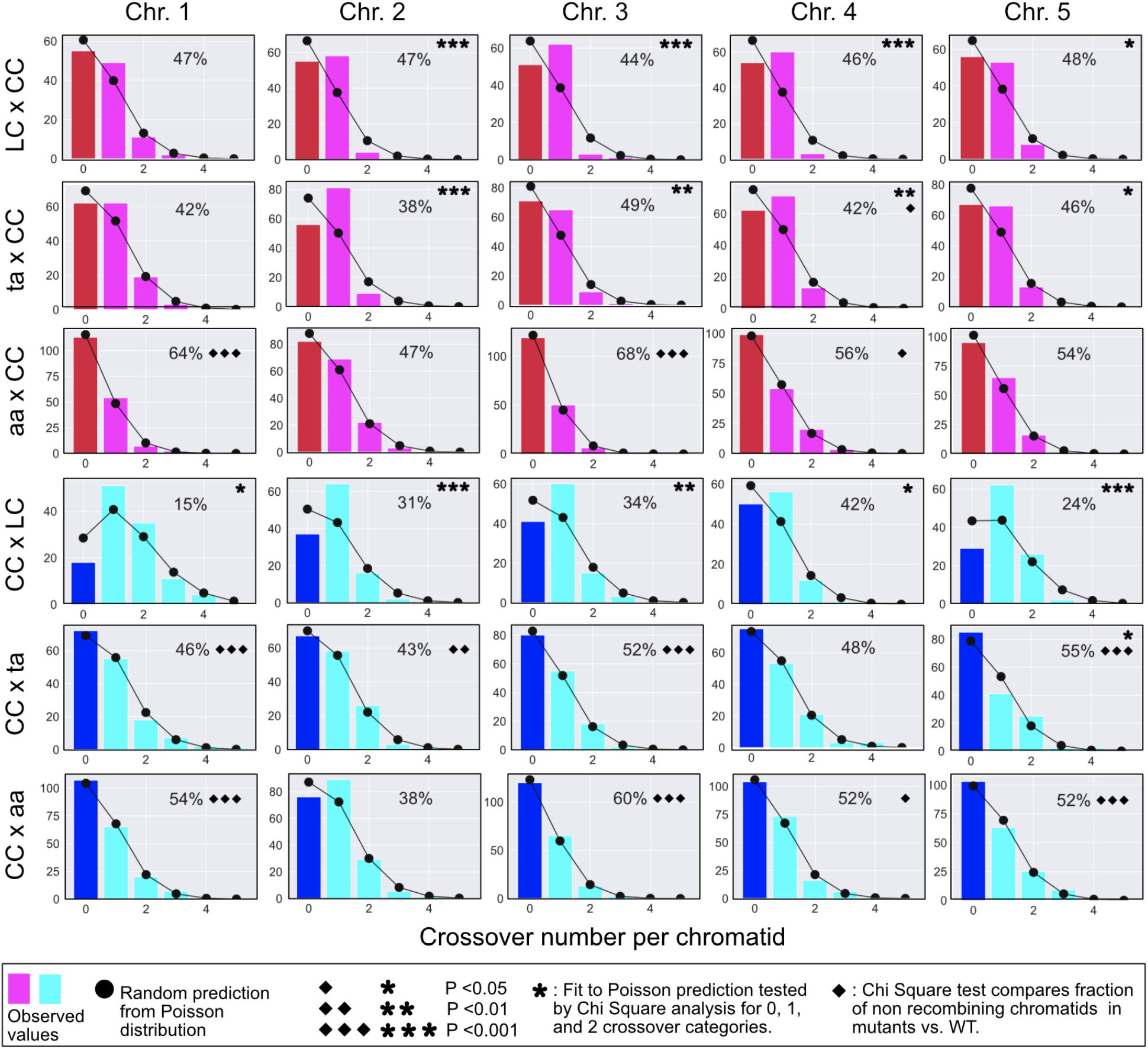
Comparison of CO numbers per chromatid in WT and *asy1* mutants. The bar plots illustrate the count of chromatid classes for given genotypes and chromosomes. The random expectation according to the Poisson distribution is displayed by the round black markers. The star markers display significance of deviation from the random fit. The percent of the 0 CO chromatid class is displayed in each plot. Significant deviation from the respective control is marked with diamonds. The exact number of instances in each category can be found in Table S3.

The hallmark of CO assurance is the presence of one obligatory CO that causes a non-random distribution of COs over the genome, resulting in a deviation from a poisson distribution of COs, and an overrepresentation of chromatids exhibiting a single CO (Fig 7, LC x CC cross). In contrast, in the ta x CC progenies, and to an even larger extent in the aa x CC progenies, we consistently observed distributions of COs that were closer to the expected random distribution (asterisks, significance test in Fig. 7). If the CO distribution deviates in the *asy1* crosses from a random distribution, it is usually due to an underrepresentation of the 1-CO class, e.g. chromosome 1 in male meiosis (Fig. 7, lower three panels), combined with an overrepresentation of both non-recombined chromatids (diamonds, significance test in Fig. 7), and chromatids with more than one CO. Thus, we conclude that another key feature of COs, i.e., CO assurance, is also affected by the loss of ASY1 function, namely, ASY1 is required for the acquisition of one obligatory CO for each homolog pair.

## Discussion

ASY1 and its homolog Hop1 are important for several aspects of meiotic recombination (8–12). Here, we have focussed on the role of ASY1 in CO placement in female and male meiosis. Mapping COs in hybrids between two different Arabidopsis accessions that are mutant for *ASY1* has allowed us to reveal several important and previously not recognized aspects of the function of ASY1, foremost that ASY1 mediates CO interference and plays an important role for CO assurance. Using two *asy1* alleles with different strengths shows that these two aspects are sensitive to the level of ASY1 activity. Our findings are consistent with another report by Lambing et al. that appeared while our paper was in revision (46). Lambing et al. demonstrate that ASY1 is involved in CO interference and counteracts telomere-led CO placements in male meiosis.

Interestingly, CO interference was also reduced in yeast cells mutant for the AAA-ATPase PACHYTENE CHECKPOINT2 (PCH2) (47). PCH2 has been previously found to be important for the removal of ASY1 and Hop1 to promote assembly of the synaptonemal complex (48–50). This raises the question of why loss of ASY1, and a failure to remove it can cause similar mutant phenotypes. However, recent work indicated that PCH2, at least in Arabidopsis, also plays a role in delivering ASY1 into the nucleus, and in the absence of PCH2 or its co-factor COMET, ASY1 strongly accumulated in the cytoplasm while ASY1 is a solely nuclear localized protein in the WT (34, 51). Thus, it is possible that the timely and correct association of ASY1 with the axis, especially during early phases of axis formation, could be compromised in *pch2* reconciling the *pch2* and *asy1* mutant effects with respect to CO interference. Conversely, we postulate that interference should also be affected in *comet* mutants.

Pairing and synapsis usually start in distal regions of homologs in Arabidopsis (52–54). Notably, despite the general failure of chromosome alignment and synapsis in the *asy1* mutant, telomere clustering and the pairing of telomeres of homologs has been found to occur during meiotic interphase and early prophase I (55). Thus, the placement of COs in the distal regions of homologs in *asy1,* as observed here and in previous studies, is consistent with this observation (12, 33). With that respect, it is interesting to note that synapsis occurs in only short chromosomal sections in Sordaria *mer2* mutants and COs tend to cluster in these sections, thus also modulating the strength of interference (56).

Remarkably, even the hypomorphic *asy1* mutant showed a strong distalization of COs. Therefore, it is plausible that even a slight modulation of the axis and/or the speed by which the axis is formed could restrict COs to distal positions while maintaining a high level of fertility as seen in the hypomorphic *asy1^T142^* mutant, which is semi fertile (34). In this context it is interesting to note that many polyploid species show a distalization of COs based on cytological analyses (29, 57). Moreover, mutations in *ASY1* and of the axis component *ASY3* have been found to be specifically associated with tetraploidy in *Arabidopsis arenosa* species, hinting at an adaptive advantage of these mutations in polyploids (31, 58). While the benefit of distalized COs in polyploids is obscure, CO numbers have often been found to be reduced in polyploids and this effect would be very beneficial for the faithful segregation of species with more than two homologs, as it would reduce the level of chromosomes that are interconnected by COs and hence prone to missegregation (29). Reduced CO number in polyploids has been speculated to be due to increased interference (29). Our findings offer an alternative explanation, i.e., reduction/slowing down of pairing with concomitant reduction of interference. Moreover, our findings would also explain why COs are distalized in polyploids. However, whether an *asy1*-dependent decrease in interference represses missegregation in polyploids can only be a valid explanation if the remaining COs that form in the distal region of chromosomes are more likely to all concern the same pair of homologs. Such an exclusive CO position could possibly be forced by sterical constraints between different homologs. Determination of CO patterns by sequencing in polyploid species will help resolve this question. In addition, live cell imaging of meiosis, as recently established for Arabidopsis (59), and tracking of chromosomes in polyploid species could be a very powerful tool to address this point.

Why interference is reduced in *asy1* mutants remains unclear. Applying the current beam-film model of interference to our data (21), leads to the hypothesis that a reduction in functional ASY1 protein levels would prevent the tension relaxation brought about by a CO to spread along the chromosome, possibly because the chromosome axis is not connected by ASY1. Consistently, the ASY1 homolog Hop1 has been proposed, based on *in vitro* data, to build long head-to-tail polymers (60, 61). Such a polymer could possibly transmit a relaxation force. However, at least in such a model, it seems unlikely that ASY1 itself contributes to the tension since, in that case, no COs would be expected in *asy1* mutants in the first place. Hence, more likely under the assumption of the beam-film model, an ASY1 polymer, if built *in vivo*, could serve as a platform for other proteins that help relaxing the mechanical stress such as topo II (62). Interestingly, TOPO II has been co-precipitated with ASY1 from *Brassica oleracea* meiocytes (63). However, it is currently not clear whether this interaction is direct or whether TOPO II in Brassica is also involved in interference control.

Our work showed that ASY1 regulates not only interference but also CO assurance. The beam-film model predicts that CO assurance is not a function of interference although the initial obligatory CO drives interference subsequently by providing the starting gun for mechanical stress. While some mutants, which affect interference in yeast, were found to still undergo an obligatory CO, such as mutants in *TOPO II*, CO assurance and interference were concomitantly compromised in others mutants, e.g. in mutants of *MutS HOMOLOG4* (*MSH4*) (19, 64). We currently do not know whether the effect of *ASY1* on interference and assurance have the same biochemical foundation, or reflect two different functions of ASY1.

Moreover, the observation that the class with one CO is often underrepresented in *asy1* mutant populations in combination with closely spaced COs in the telomeric and subtelomeric regions suggests that *asy1* mutants could represent a rare case of negative interference, i.e., the attraction of COs after a first CO is formed (5). Notably, we found that the extent of interference and assurance reduction in *asy1* mutants appears to depend on the individual chromosome for yet unknown reasons. Thus, the exploration of the dynamics and function of the chromosome axis in CO interference and CO assurance, ideally with chromosome-specific resolution, remains an exciting question.

## Methods

### Plant materials and growing conditions

The *Arabidopsis thaliana* accessions Columbia (Col-0) and Landsberg *erecta* (L*er*-1) were used as the WT references in this study. The *asy1* T-DNA insertion line (SALK_046272) in Col-0 background (65) and *mlh1-3* (SK25975) was obtained from the T-DNA mutant collection at the Salk Institute Genomics Analysis Laboratory (SIGnAL, http://signal.salk.edu/cgi-bin/tdnaexpress) via NASC (http://arabidopsis.info/). The hypomorphic non-phosphorylatable mutant *asy1^T142V^* in Col-0 background was generated previously in Yang et al. (34, 66). The *asy1* mutant in L*er* background was generated in this study by CRISPR/Cas9 approach (Fig. S1). All plants were grown in growth chambers with a 16h light/21°C and 8h dark/18°C cycle at 60% humidity. The generation of genome sequencing populations were described in the main text. For this procedure, homozygous *asy1* mutants (*asy1^Col-0^, asy1^T142V^, and asy1^ler-1^*) were used.

### Genomic sequencing and raw read processing

For Arabidopsis genomic DNA extraction, about 2 cm^2^ of leaf was harvested and used for DNA extraction using the SPRI beads-based method (67). Subsequently, genomic DNAs were quantified using a plate fluorometer with SYBR Green I, and normalized. After normalization, the samples were checked on agarose for uniformity. The generation of the sequencing libraries was performed following the Rowan protocol (68). Briefly, samples were fragmented, adapters were added, the fragments were amplified with indexed P5 and P7 primers, and visualization on agarose before being pooled (see Supplemental Dataset 2 for index and pooling information). Pools then underwent size selection with magnetic beads, before measuring DNA concentration using a Qubit (Invitrogen) and verifying size distribution with the Agilent Bioanalyzer. Pools were sequenced on an Illumina HiSeq 4000 sequencer, to obtain PE150 reads. Raw reads were checked for sequence quality, trimmed, and demultiplexed using custom python scripts, as described previously (69). Next, they were aligned to the TAIR10 genomic reference (https://www.arabidopsis.org) using BWA and default parameters (70), producing sam files appropriate for dosage analysis (see below). For the purpose of genotyping and the creation of genetic maps, the data was also concatenated into a single mpileup.txt file using samtools (71). Finally, this mpileup file was further processed to obtain the percentages of each base call at each position and each sample using the script called mpileup-parser.py (see http://comailab.genomecenter.ucdavis.edu/index.php/Mpileup for download and documentation).

### Dosage analysis and genotyping

Aneuploidy and dosage variation detection was performed as previously described (72). Briefly, the genome was divided into consecutive non-overlapping 100 kb bins and read that mapped to each bin were counted for each sample using a custom python script (bin-by-sam.py, available on the Comai Laboratory website: http://comailab.genomecenter.ucdavis.edu/index.php/Main_Page). For each bin, read counts were normalized to account for variation in total read counts, as well as compared to the mean values for the whole population, in order to detect variation in dosage relative to the expected value of 2 in a diploid background. Data were plotted (Fig. 1 for example) and aneuploidy was detected visually, as a variation in copy number that is consistent across at least 3 consecutive bins.

### Determination of CO locations and data analysis

A list of Col-0/Ler-1 SNPs was derived from a previously published list of ∼870K SNP positions, (http://mtweb.cs.ucl.ac.uk/mus/www/19genomes/variants.SDI/ler_0.v7c.sdi (73)). We first filtered this list by selecting positions that had been detected by both the IMR and DENOM software, that did not include any ambiguous bases (only A, T, G and C were allowed), with phred scores > 100, and for which the reported consensus base SNP and the high Mapping Quality consensus base SNP were in agreement. This reduced the SNP set to ∼418K positions. Next, we further filtered this list by only retaining positions that were covered at least once in at least 500/1108 of our samples and for which two both expected alleles were detected in our population (at least 5% of each allele type). The resulting ∼125K positions were used for genotyping. For each sample, the alleles calls at each of these SNP positions was recorded. Because we sequenced each sample at low coverage, allele calls were pooled in order to obtain robust and complete genotype calls for each sample. Specifically, allele calls were pooled into a total of 239 consecutive non-overlapping 500 kb bins, and a per bin genotype was derived for each individuals by calculating the mean percentage of Col-0 allele in each bin, as previously described (74). The resulting genotype information was input into R/qtl for the creation of genetic maps for each cross (est.map function), and the detection of COs (countXO function) and the determination of their position (locateXO function) (75).

### CO interference

The testcross design described above was used to assess interference. A Python script was used to divide the genome into non-overlapping intervals of arbitrary length. Recorded recombination breakpoints were counted inferring COs per interval. The recombination frequency of any interval was equal to the (number of COs)/(number of scored chromatids). Predicted double COs number was determined by taking the square of the recombination rate. Double COs instances were scored based on COs pairs. Ten instances of triple or higher number CO’s were observed in the 4.5Mb bin analysis, and scored as individual instances. For example, an interval with three COs would be scored as having 2 double COs, one between CO1 and CO2, the second between CO2 and CO3. These data are summarized in Supplemental Dataset 3.

### Cytologenetic analyses

Meiotic chromosome spreads and fluorescence *in situ* hybridization (FiSH) were performed according to procedures previously described (76). The following DNA probes were used: pTa71 [45S ribosomal DNA (rDNA), pTa71 of *Triticum aestivum*; (77)] and pCT4.2 [5S 133 rDNA; (78)].

Immunolocalization procedures were conducted following the method detailed in Armstrong *et al.* (55), with modifications detailed in Varas and Pradillo (79). The primary antibodies used were kindly provided by Professor Chris Franklin (University of Birmingham): anti-AtASY1 (rat; 1/1000), anti-AtZYP1 (rat; 1/500), anti-MLH1 (rabbit, 1/500), anti-RAD51 (rabbit, 1/300). Secondary antibodies were anti-rat and anti-rabbit IgG FITC conjugated (Agrisera) and anti-rabbit and anti-rat IgG Alexa Fluor 555 conjugated (Molecular Probes).

Slides were examined on an Olympus BX61 epifluorescence microscope and imaged with a CCD Olympus DP71 camera and computer using analySIS software (Soft Imaging System). For foci scoring blind-coded digital images were used. Adobe Photoshop (CC 2018 v.19.0) and ImageJ (version 1.51s) were applied for image processing.

## Accession Numbers

Sequence data from this article can be found in the NCBI SRA database under accession number TBA.

## Author Contributions

A.S., I.H. and Lu..C. designed the research; G.P., I.M., C.Y., N.F.J., N.L., L.C., H.T.T., M.P., performed the experiments. G.P., I.M., C.Y., M.P., Lu.C., A.S. analysed the data. I.H., C.Y., M.P., Lu.C., A.S. wrote the manuscript.

## Acknowledgments

We thank Beth Rowan for assistance with the preparation of the DNA sequencing libraries, and Meric Lieberman for bioinformatics assistance. We are grateful to Kirsten Bomblies (ETH Zurich), Nancy Hollingsworth (Stony Brook University, New York), and Erik Wijnker (University of Wageningen) and for critical reading and helpful comments on the manuscript. The ASY1 and MLH1 primary antibodies were kindly provided by Chris Franklin (University of Birmingham, United Kingdom). NFJ is a PhD fellow funded by the FPU programme of Spanish Ministry of Education (FPU16/02772). This research was supported by the European Union (Marie Curie ITN, COMREC 606956 and MEICOM 765212) to M.P. and A.S. M.P. and A.S. are members of the COST Action n° CA 16212 ‘In Depth’ http://www.cost.eu/COST_Actions/ca/CA16212. MP acknowledges the support of the Ministry of Economy and Competitiveness of Spain (AGL2015-67349-P). MP is part of the International Plant Nucleus Consortium (IPNC, https://radar.brookes.ac.uk). L.C. and I.M.H. are supported by the Innovative Genomics Institute. C.Y. is supported by the Start-up fund provided by Huazhong Agricultural University. A.S. kindly acknowledges core funding of the University of Hamburg.

## Supplemental information

**Table S1.** Frequency of univalents in ta, aa and *asy1-4* (Col-0) mutants.

| Mutants | Number of univalents | Number of cells | Percentage |
| --- | --- | --- | --- |
| *ta* | 8 | 1 | 2.44% |
|  | 6 | 1 | 2.44% |
|  | 4 | 14 | 34.15% |
|  | 2 | 14 | 34.15% |
|  | 0 | 11 | 26.83% |
| *aa* | 8 | 8 | 16.00% |
|  | 6 | 26 | 52.00% |
|  | 4 | 14 | 28.00% |
|  | 2 | 2 | 4.00% |
| *asy1-4* (Col-0) | 10 | 3 | 6.38% |
|  | 8 | 6 | 12.77% |
|  | 6 | 19 | 40.43% |
|  | 4 | 14 | 29.79% |
|  | 2 | 5 | 10.64% |
|  | 0 | 0 | 0% |

**Table S2.**
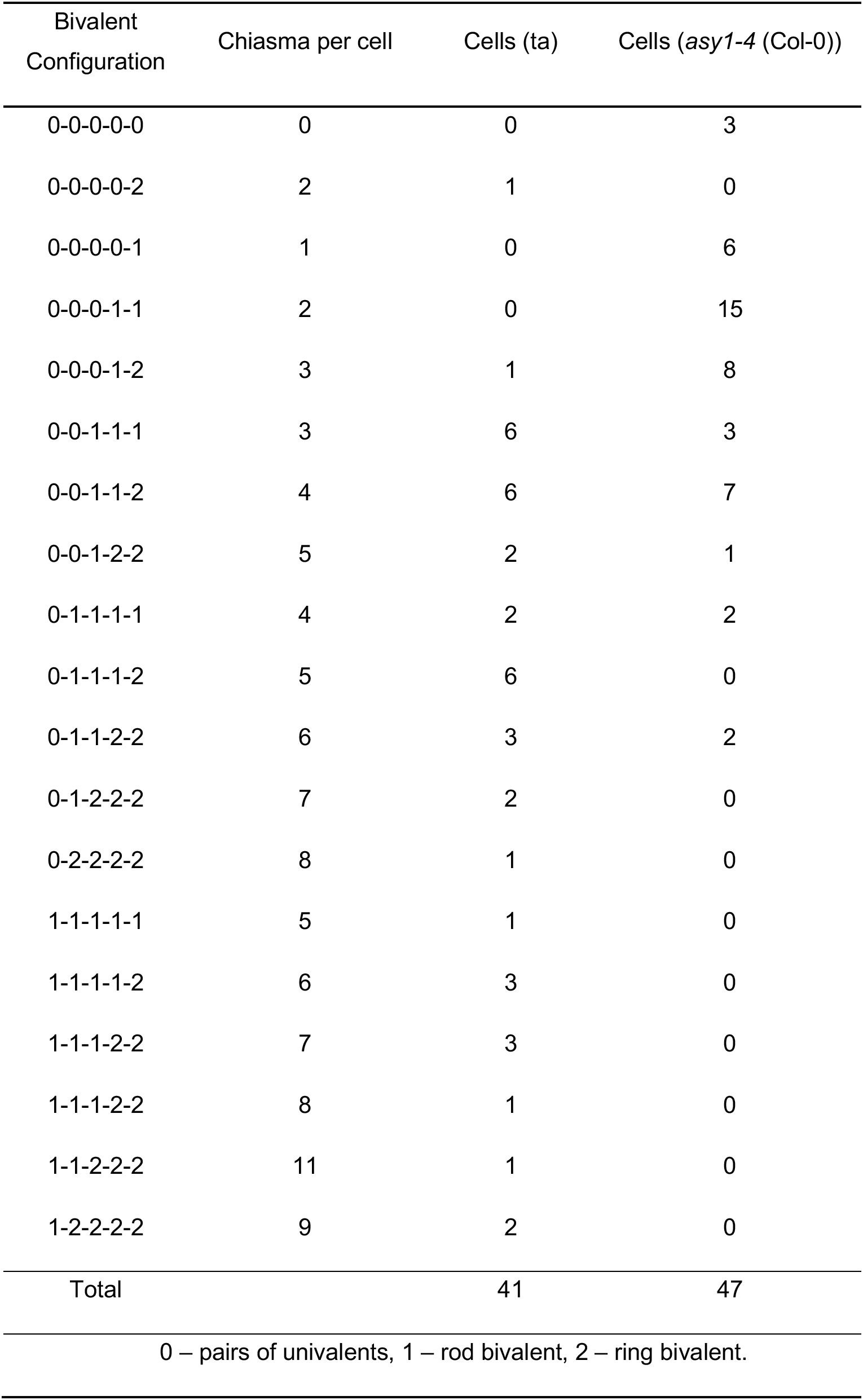
Chromosome configurations in ta and *asy1-4* (Col-0) mutants.

| Bivalent Configuration | Chiasma per cell | Cells (*ta*) | Cells (*asy1-4* (Col-0)) |
| --- | --- | --- | --- |
| 0-0-0-0-0 | 0 | 0 | 3 |
| 0-0-0-0-2 | 2 | 1 | 0 |
| 0-0-0-0-1 | 1 | 0 | 6 |
| 0-0-0-1-1 | 2 | 0 | 15 |
| 0-0-0-1-2 | 3 | 1 | 8 |
| 0-0-1-1-1 | 3 | 6 | 3 |
| 0-0-1-1-2 | 4 | 6 | 7 |
| 0-0-1-2-2 | 5 | 2 | 1 |
| 0-1-1-1-1 | 4 | 2 | 2 |
| 0-1-1-1-2 | 5 | 6 | 0 |
| 0-1-1-2-2 | 6 | 3 | 2 |
| 0-1-2-2-2 | 7 | 2 | 0 |
| 0-2-2-2-2 | 8 | 1 | 0 |
| 1-1-1-1-1 | 5 | 1 | 0 |
| 1-1-1-1-2 | 6 | 3 | 0 |
| 1-1-1-2-2 | 7 | 3 | 0 |
| 1-1-1-2-2 | 8 | 1 | 0 |
| 1-1-2-2-2 | 11 | 1 | 0 |
| 1-2-2-2-2 | 9 | 2 | 0 |
| Total |  | 41 | 47 |
| 0 – pairs of univalents, 1 – rod bivalent, 2 – ring bivalent. |  |  |  |

**Table S3.** CO number and frequency per chromatid in WT and *asy1* mutants.

| # CO per chromatid |  | Chromosome 1 |  |  |  |  |  | Chromosome 2 |  |  |  |  |  | Chromosome 3 |  |  |  |  | Chromosome 4 |  |  |  |  |  | Chromosome 5 |  |  |  |  |
| --- | --- | --- | --- | --- | --- | --- | --- | --- | --- | --- | --- | --- | --- | --- | --- | --- | --- | --- | --- | --- | --- | --- | --- | --- | --- | --- | --- | --- | --- |
|  |  | 0 | 1 | 2 | 3 | 4 | 5 | 0 | 1 | 2 | 3 | 4 |  | 0 | 1 | 2 | 3 |  | 0 | 1 | 2 | 3 | 4 |  | 0 | 1 | 2 | 3 | 4 |
| #<br>Individuals | Cross Type |  |  |  |  |  |  |  |  |  |  |  |  |  |  |  |  |  |  |  |  |  |  |  |  |  |  |  |  |
|  | LC x CC | 55 | 49 | 11 | 2 | 0 | 0 | 55 | 58 | 4 | 0 | 0 |  | 51 | 62 | 3 | 1 |  | 54 | 60 | 3 | 0 | 0 |  | 56 | 53 | 8 | 0 | 0 |
|  | ta x CC | 62 | 62 | 19 | 3 | 0 | 0 | 56 | 81 | 9 | 0 | 0 |  | 71 | 65 | 9 | 1 |  | 62 | 71 | 13 | 0 | 0 |  | 67 | 66 | 13 | 0 | 0 |
|  | aa x CC | 113 | 54 | 7 | 2 | 0 | 0 | 82 | 69 | 22 | 3 | 0 |  | 119 | 50 | 6 | 1 |  | 99 | 54 | 20 | 3 | 0 |  | 95 | 65 | 16 | 0 | 0 |
|  | CC x LC | 18 | 51 | 35 | 11 | 4 | 0 | 37 | 64 | 16 | 2 | 0 |  | 41 | 60 | 15 | 3 |  | 50 | 56 | 12 | 1 | 0 |  | 29 | 62 | 26 | 2 | 0 |
|  | CC x ta | 72 | 55 | 18 | 7 | 2 | 1 | 67 | 58 | 26 | 3 | 1 |  | 80 | 55 | 18 | 2 |  | 75 | 53 | 21 | 3 | 3 |  | 85 | 41 | 25 | 2 | 2 |
|  | CC x aa | 107 | 65 | 20 | 7 | 1 | 0 | 76 | 89 | 29 | 5 | 1 |  | 120 | 65 | 13 | 2 |  | 104 | 73 | 16 | 6 | 1 |  | 103 | 63 | 25 | 9 | 0 |
| %<br>Individuals | Cross Type |  |  |  |  |  |  |  |  |  |  |  |  |  |  |  |  |  |  |  |  |  |  |  |  |  |  |  |  |
|  | LC x CC | 47 | 42 | 9 | 2 | 0 | 0 | 47 | 50 | 3 | 0 | 0 |  | 44 | 53 | 3 | 1 |  | 46 | 51 | 3 | 0 | 0 |  | 48 | 45 | 7 | 0 | 0 |
|  | ta x CC | 42 | 42 | 13 | 2 | 0 | 0 | 38 | 55 | 6 | 0 | 0 |  | 49 | 45 | 6 | 1 |  | 42 | 49 | 9 | 0 | 0 |  | 46 | 45 | 9 | 0 | 0 |
|  | aa x CC | 64 | 31 | 4 | 1 | 0 | 0 | 47 | 39 | 13 | 2 | 0 |  | 68 | 28 | 3 | 1 |  | 56 | 31 | 11 | 2 | 0 |  | 54 | 37 | 9 | 0 | 0 |
|  | CC x LC | 15 | 43 | 29 | 9 | 3 | 0 | 31 | 54 | 13 | 2 | 0 |  | 34 | 50 | 13 | 3 |  | 42 | 47 | 10 | 1 | 0 |  | 24 | 52 | 22 | 2 | 0 |
|  | CC x ta | 46 | 35 | 12 | 5 | 1 | 1 | 43 | 37 | 17 | 2 | 1 |  | 52 | 35 | 12 | 1 |  | 48 | 34 | 14 | 2 | 2 |  | 55 | 26 | 16 | 1 | 1 |
|  | CC x aa | 54 | 33 | 10 | 4 | 1 | 0 | 38 | 45 | 15 | 3 | 1 |  | 60 | 33 | 7 | 1 |  | 52 | 37 | 8 | 3 | 1 |  | 52 | 32 | 13 | 5 | 0 |

C: Col-0, t: *asy1*<sup>T142V-Col-0</sup>, a: *asy1*<sup>Ler-1</sup> or *asy1*<sup>Col-0</sup> null alleles. Control crosses are highlighted in grey.

**Supplemental Fig. S1.**
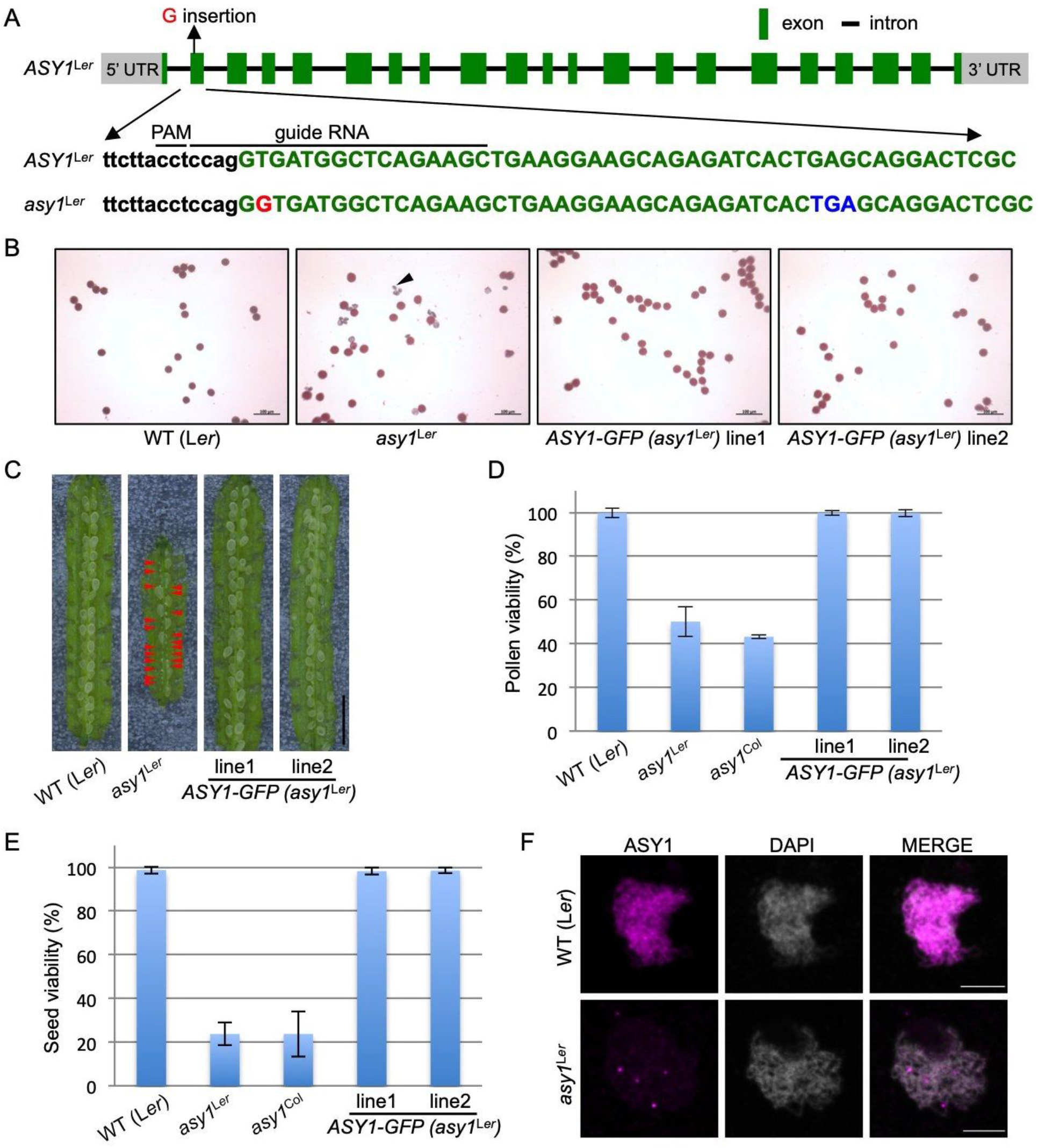
Phenotypic characterization of *asy1* mutants generated by CRISPR/Cas9 in the Arabidopsis accession Landsberg *erecta* (L*er-*1) (A) Map of the *ASY1* genomic region in L*er*-1. The targeting sequence (guide RNA) and PAM sequence are indicated by black lines. A G-nucleotide insertion in the mutant is highlighted in red and the resulting premature stop codon in blue. (B) Peterson staining of pollens in L*er-*1 WT (WT), *asy1^Ler-1^* mutants, and two *ASY1-GFP* (*asy1^Ler-1^*) complementation lines. Blue staining indicates dead pollen (arrowhead). Scale bar: 100 μm. (C) Seed sets in silique of WT, *asy1^Ler-1^*, and *ASY1-GFP* (*asy1^Ler-1^*) complementation lines. Red arrowheads indicate aborted seeds. Scale bar: 2 mm. (D) Quantification of pollen viability in WT, *asy1^Ler-1^*, *asy1^Col-0^*, and *ASY1-GFP* (*asy1^Ler-1^*) complementation lines. (E) Quantification of seed viability in in WT, *asy1^Le-1r^*, *asy1^Co-0l^*, and *ASY1-GFP* (*asy1^Ler-1^*) complementation lines. Error bars represent standard deviations. (F) Immuno localization of ASY1 in L*er*-1 WT plants in L*er*-1 *asy1* mutants. While ASY1 is strongly associated with the chromosome axis in the WT, ASY1 could not be detected in *asy1* mutant plants, indicating that the generated CRISPR/cas9 mutant is an *asy1* null allele. Size bar is 5µm.

**Supplemental Fig. S2.**
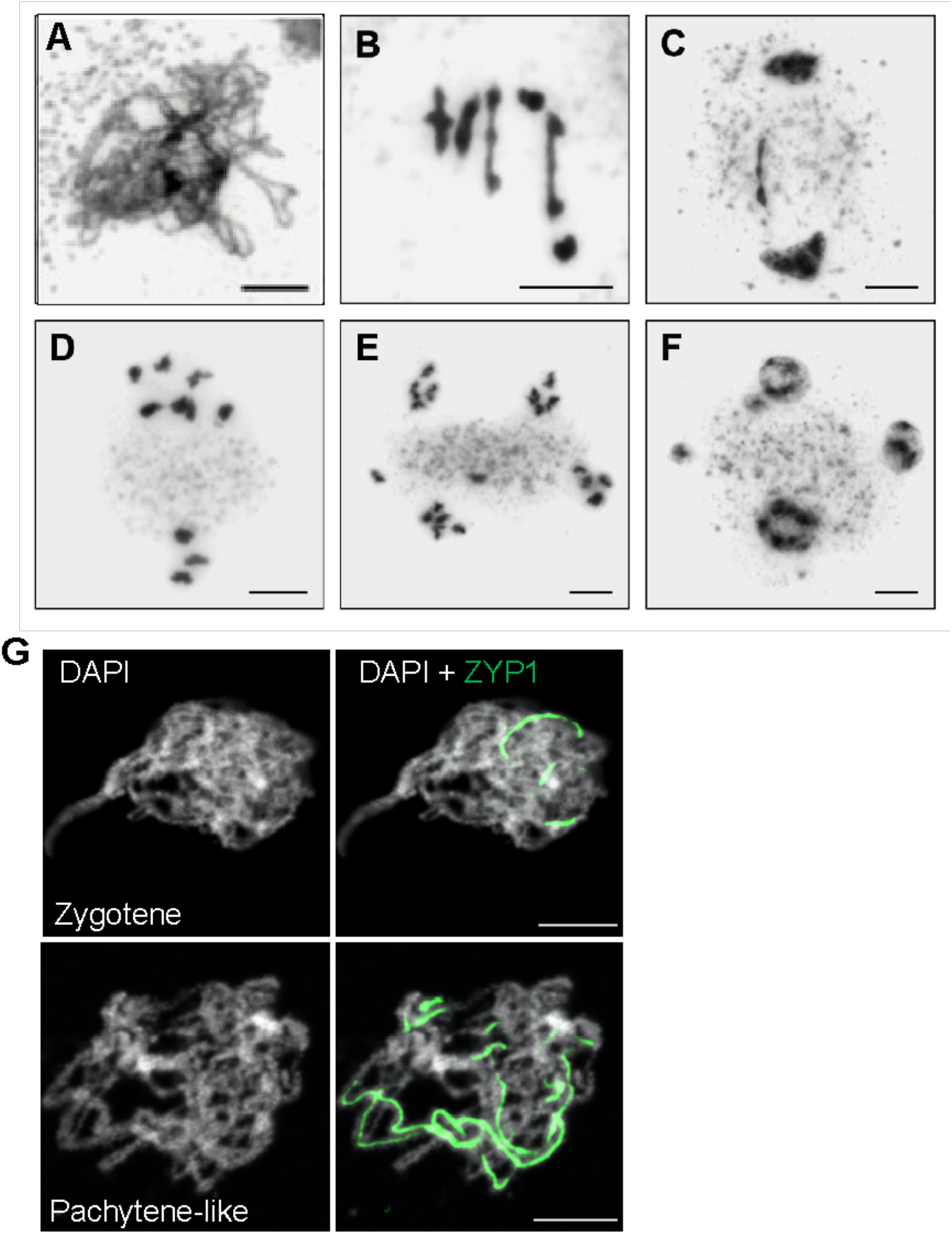
Cytology of male meiosis in ta plants. (A) Zygotene. The ta mutant is asynaptic, showing failure of synapsis during prophase I. (B) Metaphase I with two univalents. We observed univalents in more than 70% (29/41) of the metaphases I analyzed. The proportion of cells with different number of univalent was shown in Supplemental table 1. (C) Anaphase I with a bridge. (D) Metaphase II with unbalanced nuclei (7:3). (E) Anaphase II displaying unequal chromosome segregation. Two chromosomes are located at the organelle band. (F) Polyad with two micronuclei. (G) Immunolocalization of ZYP1 shows the defective synapsis in ta mutant. Bars represent 5µm.

**Supplemental Fig. S3.**
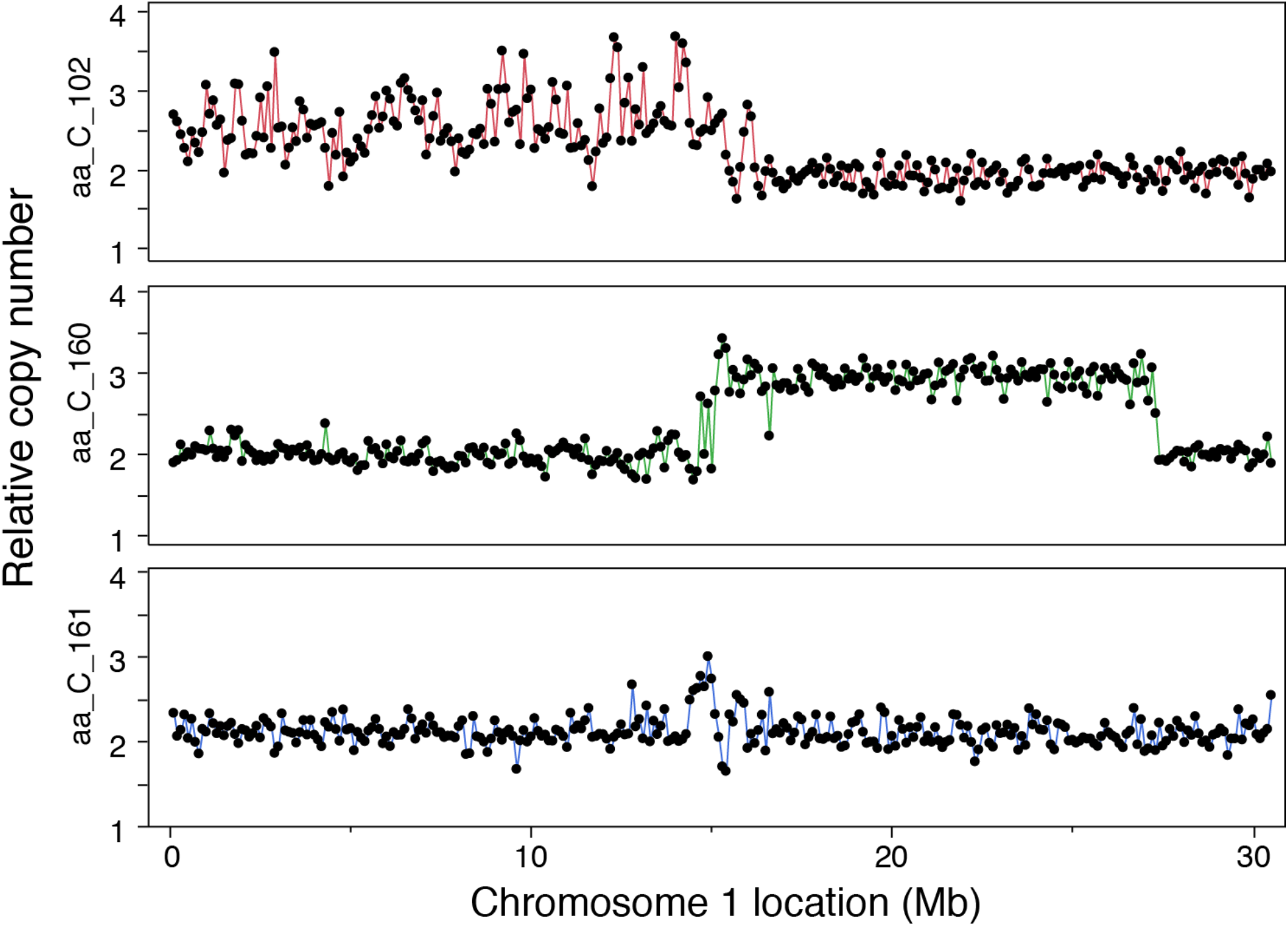
Extreme genome instability in the progeny of the *asy1* null mutant. For each individual, relative dosage is shown based on sequence coverage variation after normalization to the whole population. Dots represents non-overlapping consecutive 100kb bin. One of the progenies from the aa x CC cross carries a third copy of the top arm of chromosome 1, which exhibits widespread dosage variation (#102, top panel). In comparison, variation is much less pronounced in a control diploid individual (#161, bottom panel), or in another aneuploid individual carrying a third copy of a middle segment of chromosome 1 (#160, middle panel).

**Supplemental Fig. S4.**
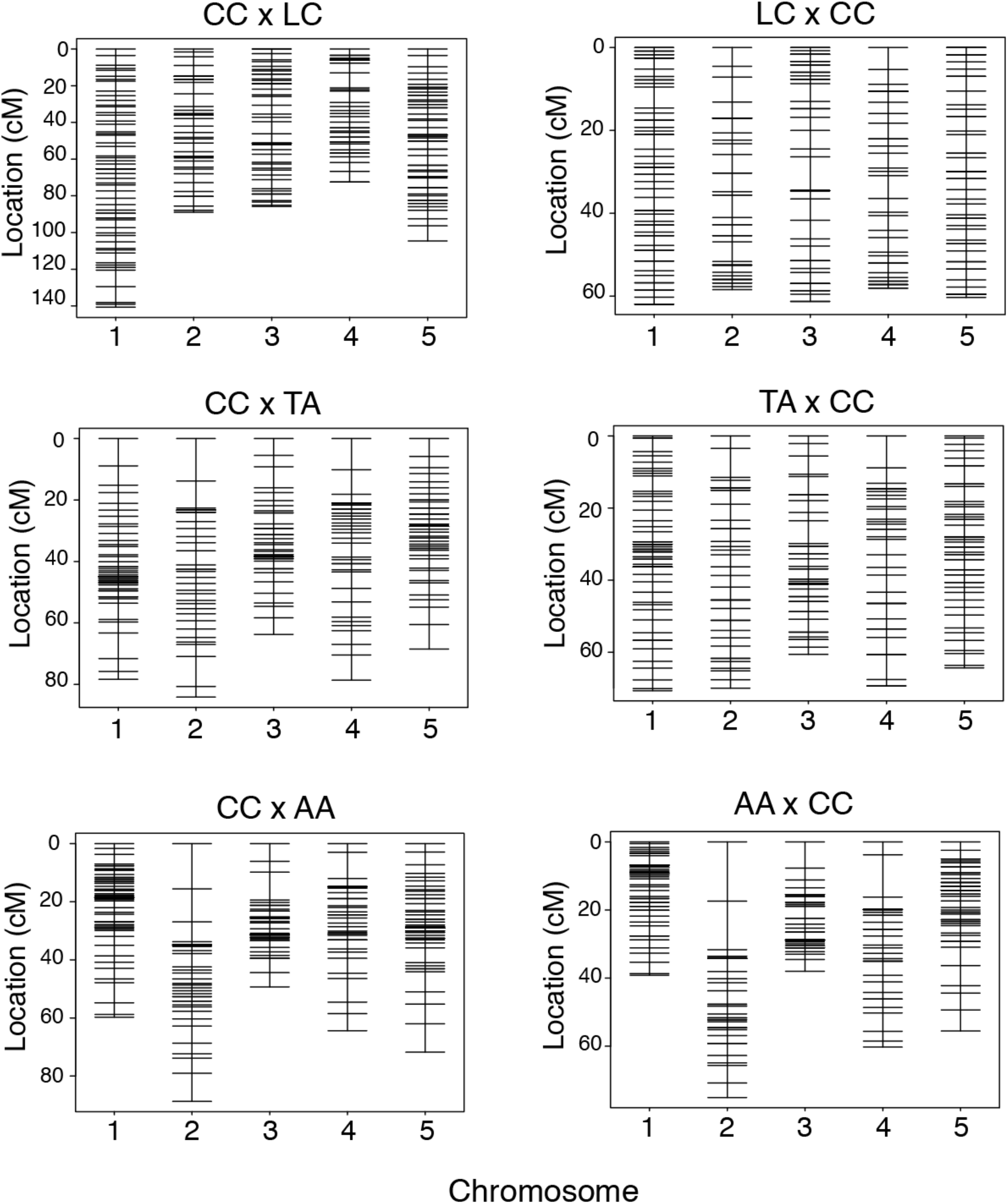
Genetic maps of the 6 populations analyzed. For each population, 239 markers were analyzed.

**Supplemental Fig. S5.**
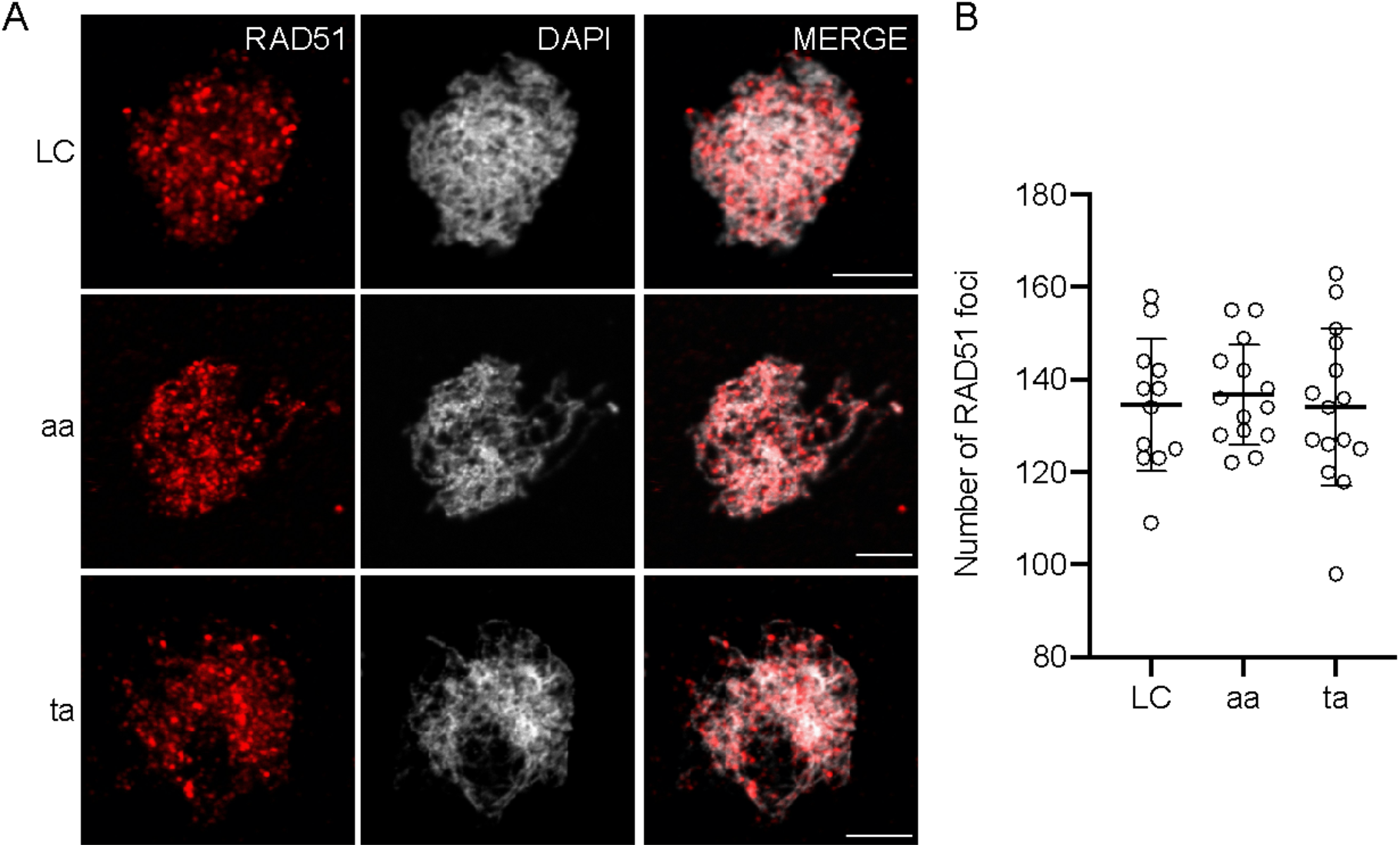
Detection of DSBs via analysis of RAD51 localization. (A) Early stages of prophase I (leptotene/zygotene) after immunolabelling to detect the DSB marker RAD51 (red). DAPI staining appears in grey. Bars represent 5µm. (B) Analysis of RAD51 foci showing that the number of DSBs are neither reduced in ta nor in aa plants, in comparison to the WT LC plants. Each dot represents an individual cell and bars indicate the mean.

**Supplemental Fig. S6.**
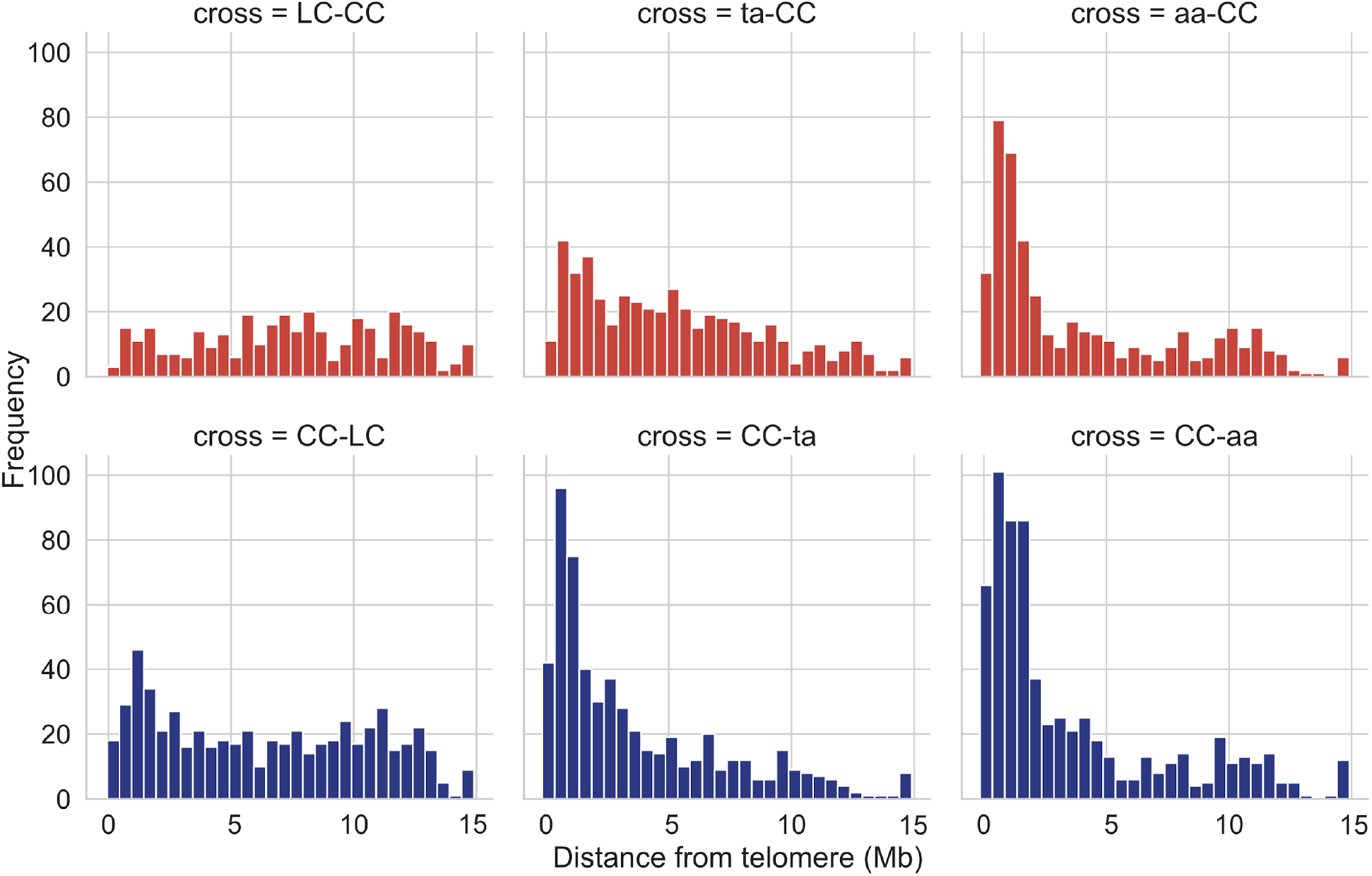
Distribution of CO location for each cross. The histograms display the distribution of distances between each CO and the nearest telomere. Data aggregate all chromosomes and arms. Knock out of *ASY1* is associated with a higher proportion of COs close to the telomeres. The hypomorphic mutants display in intermediate distribution.

**Supplemental Fig. S7.**
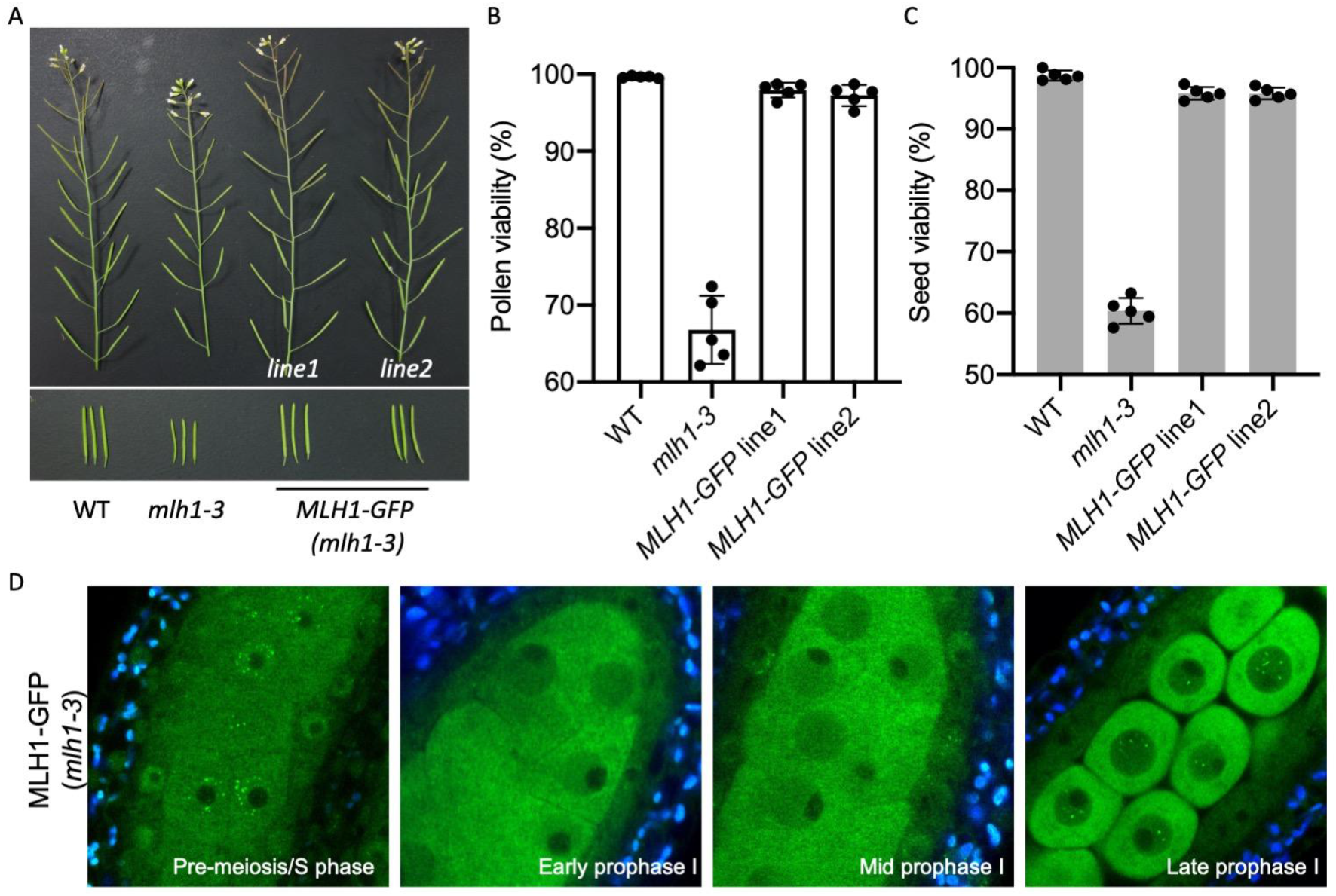
MLH1-GFP complements the meiotic defects in *mlh1* mutant and forms foci in pre-meiosis/S phase and late prophase I stages. (A) Main branches (upper panel) and siliques (lower panel) of the wildtype (WT), *mlh1*, and two reference lines expressing the MLH1-GFP construct in the *mlh1* mutant background. (B) Pollen viabilities in WT, *mlh1*, and the two reference lines expressing MLH1-GFP (in *mlh1-3)*. (C) Seed viabilities in WT, *mlh1*, and the two reference lines expressing MLH1-GFP (in *mlh1-3)*. (D) Expression and localization of MLH1-GFP in male meiocytes of *mlh1* mutant plants at different meiotic stages.

**Supplemental Fig. S8.**
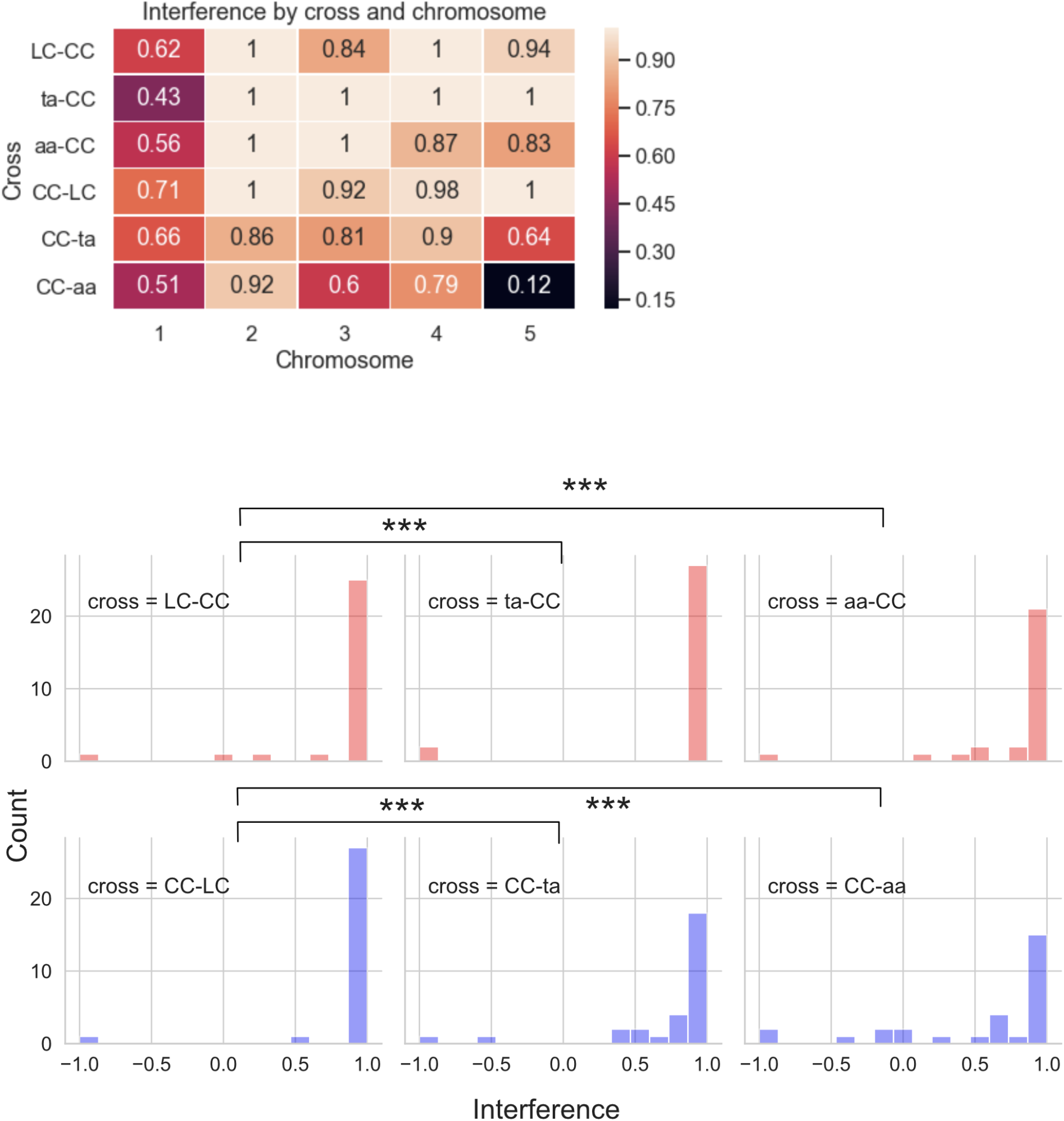
Interference analysis. Interference was measured in a sliding 4.5 Mb window. Top: mean interference per chromosome and cross. Bottom: Distribution of interference value in the 30 bins sampled for each cross (135/4.5 Mb). For display purposes, extreme interference values were clipped at −1 and +1. Knock out of *ASY1* is associated with reduced interference. P-value: *** < 0.001 according t.

## Supplemental Datasets

**Supplemental Dataset 1: List of COs detected in the 6 populations. The cross-over positions are indicated in Mb.**

**Supplemental Dataset 2: List of samples and barcodes used for demultiplexing after Illumina sequencing.**

**Supplemental Dataset 3: Interference analysis using 4.5 Mb windows.**

## Supplemental Movies

**Supplemental Movie 1. Z-tacks of MLH1-GFP in male meiocytes of wildtype plant.**

**Supplemental Movie 2. Z-tacks of MLH1-GFP in male meiocytes of *asy1* mutant.**

**Supplemental Movie 3. Z-tacks of MLH1-GFP in male meiocytes of *asy1* mutant.**

**Supplemental Movie 4. Z-tacks of MLH1-GFP in male meiocytes of *asy1* mutant.**

## Notes

### Competing Interest Statement

The authors have declared no competing interest.

